# Influenza A virus (H1N1) infection induces microglia activation and temporal dysbalance in glutamatergic synaptic transmission

**DOI:** 10.1101/2021.08.30.458184

**Authors:** Henning Peter Düsedau, Johannes Steffen, Caio Andreeta Figueiredo, Julia Désirée Boehme, Kristin Schultz, Christian Erck, Martin Korte, Heidi Faber-Zuschratter, Karl-Heinz Smalla, Daniela Dieterich, Andrea Kröger, Dunja Bruder, Ildiko Rita Dunay

## Abstract

Influenza A virus (IAV) causes respiratory tract disease and is responsible for seasonal and reoccurring epidemics affecting all age groups. Next to typical disease symptoms such as fever and fatigue, IAV infection has been associated with behavioral alterations presumably contributing to the development of major depression. Previous experiments using IAV/H1N1 infection models have shown impaired hippocampal neuronal morphology and cognitive abilities, but the underlying pathways have not been fully described. In this study, we demonstrate that infection with a low dose non-neurotrophic H1N1 strain of IAV causes ample peripheral immune response followed by a temporary blood-brain-barrier disturbance. Although histological examination did not reveal obvious pathological processes in the brains of IAV-infected mice, detailed multidimensional flow cytometric characterization of immune cells uncovered subtle alterations in the activation status of microglia cells. More specifically, we detected an altered expression pattern of major histocompatibility complex class I and II, CD80, and F4/80 accompanied by elevated mRNA levels of CD36, CD68, C1QA, and C3, suggesting evolved synaptic pruning. To closer evaluate how these profound changes affect synaptic balance, we established a highly sensitive multiplex flow cytometry-based approach called Flow Synaptometry. The introduction of this novel technique enabled us to simultaneously quantify the abundance of pre- and postsynapses from distinct brain regions. Our data reveal a significant reduction of VGLUT1 in excitatory presynaptic terminals in the Cortex and Hippocampus, identifying a subtle dysbalance in glutamatergic synapse transmission upon H1N1 infection in mice. In conclusion, our results highlight the consequences of systemic IAV-triggered inflammation on the central nervous system and the induction and progression of neuronal alterations.

## 5. Introduction

Influenza A virus (IAV) causes infection of the respiratory tract and repeatedly appears throughout the lifetime of almost every individual. Subtypes of IAV, such as H1N1, often contribute to seasonal epidemics or may even cause pandemics, thus representing a major cause for morbidity and mortality worldwide [1]. Approximately 8% of the U.S. population become infected with IAV each season [2], eventually resulting in over 30,000 IAV-associated deaths per year [3]. Based on these assumptions, it has been extrapolated that influenza infections sum up to 4-50 million annual cases and 15-70,000 deaths in the EU [4, 5]. The resolution of IAV infection requires the activation of innate and adaptive immunity and is initiated in lung epithelial cells, which are primary targets of IAV. Here, IAV is detected by cytosolic pattern recognition receptors such as endosomal toll-like receptors or retinoic acid inducible gene I, resulting in the initiation of NF-κB-dependent release of a plethora of cytokines and chemokines including interleukin (IL)-1β, IL-6, tumor necrosis factor (TNF), and type I interferons (IFN) that induce an anti-viral state via IFNAR1 and IFNAR2 receptor activation. The induction of interferon-regulated pathways is characterized by the expression of various GTPases such as MX proteins that interfere with virus growth in infected cells [6]. Potent activation of innate immunity results in the subsequent establishment of adaptive cellular immunity mediated by virus-specific CD8^+^ cytotoxic T cells and CD4^+^ T helper cells, and humoral immunity via antibody-producing B cells [7]. As a consequence of the mounted immune response, IAV infection is associated with high serum levels of pro-inflammatory cytokines resulting in distinct behavioral alterations in patients, a phenomenon also referred to as sickness behavior [8]. Although the profound mechanism for the induction of sickness behavior is not fully understood, previous reports have highlighted that this effect is caused by a transport of peripheral cytokines into the brain parenchyma via the blood-brain barrier (BBB) [9, 10]. Thus, it has been speculated that infections such as IAV pneumonia may act as one of the initial triggers contributing to the development of neurological disorders and therefore have been studied in experimental models over the past years [11–14].

As brain-resident macrophages, microglia cells represent the first line of defense against pathogens. Under homeostatic conditions, microglia cells possess a highly ramified morphology, constantly scanning their environment for pathogens as well as supporting neuronal function and synaptic plasticity [15–18]. Microglia become activated upon sensing of pathogens and inflammatory signals, releasing cytokines, chemokines, and other inflammatory mediators such as nitric oxide and complement factors [19]. Therefore, microglia activation has often been linked to neuropathologies like Alzheimer’s disease (AD) or schizophrenia and moreover, to aggravated phagocytosis of synapses [20, 21]. Although previous reports indicate that peripheral inflammation induced by viral infection or polyinosinic:polycytidylic acid (poly[I:C]) and lipopolysaccharides (LPS) administration leads to microglia activation and behavioral changes [22–24], the underlying pathways of neurological alterations have not been fully understood. Here we demonstrate that respiratory IAV/PR8/H1N1 infection results in a distinct peripheral inflammatory pattern that is associated with transient dysbalance in BBB homeostasis. Even though initial histological examination of brain sections did not reveal evidence for pathological changes, detailed flow cytometric characterization highlighted distinct region-specific activation of microglia in the early phase of infection. We detected that expression of markers associated with synaptic pruning peak at 14 days post infection (dpi) alongside with diminished mRNA levels of the presynaptic glutamate transporter VGLUT1. Notably, the establishment of a novel flow cytometry-based approach (“Flow Synaptometry”) enabled us to assess the composition of excitatory and inhibitory synapses, demonstrating a significant reduction of VGLUT1 in excitatory presynapses at 21 dpi, which is further accompanied by changes in neurotrophin gene expression. Our findings provide detailed insights into temporal effects of peripheral inflammation on brain homeostasis and microglia-neuronal interaction that specifically shape synapse physiology over the course of IAV infection, possibly contributing to persisting neurological alterations.

## 6. Material and Methods

### Animals

The experiments were performed with female C57BL/6JRj (8 weeks old, purchased from Janvier, Cedex, France). Animals were group-housed under specific pathogen-free conditions in individual-ventilated cages with a 12 h day/night cycle with free access to food and water. All animal care was in accordance with institutional guidelines and experiments were performed in accordance to German National Guidelines for the Use of Experimental Animals and the protocol was approved by the Landesverwaltungsamt Saxony-Anhalt.

### IAV infection

The virus stock of mouse-adapted IAV PR/8/A/34(H1N1) was derived from Madin-Darby Canine Kidney cells as described previously [25]. First, mice were anesthetized by intraperitoneal injection of ketamine (1 %)-xylazine (10 mg/ml) and ointment was applied to keep eyes hydrated. Subsequently mice were intranasally (i.n.) inoculated with 0.04 of the median lethal dose (MLD)_50_ of IAV diluted in PBS whereas mock-infected animals received PBS only [26].

### Isolation of bronchoalveolar lavage fluid, tissue, and cells

To collect bronchoalveolar lavage fluid (BAL fluid), the trachea was punctured and a vein catheter was carefully inserted. BAL was performed using 1 ml ice-cold phosphate-buffered saline (PBS). BAL fluid was spun down at 420 x *g* for 10 min. Aliquots of BAL supernatants were stored at -80 °C. Tissue collection and cell isolation was performed as described previously [27]. In brief, mice were deeply anesthetized with Isoflurane (CP Pharma, Burgdorf, Germany) and perfused intracardially with 60 ml sterile PBS prior brain extraction. For subsequent analysis by flow cytometry, brains were dissected into specific regions (Cortex - CTX, Hippocampal formation – HPF) according to the Allen Mouse Brain Atlas [28], collected in separate tubes and homogenized in a buffer containing Hank’s balanced salt solution (HBSS, Gibco, New York, NY, USA), 13 mM HEPES (pH 7.3, Thermo Fisher Scientific, Waltham, MA, USA) and 0.68 % glucose before sieving through a 70 µm cell strainer. The homogenate was fractioned on a discontinuous 30-70 % Percoll gradient (GE Healthcare, Chicago, IL, USA). Immune cells were collected from the 30/70 % Percoll interphase, washed in PBS and immediately processed for subsequent flow cytometric analysis. For RNA isolation or synaptosome preparation, perfused brains were dissected as described above and either stored in RNAlater solution (Thermo Fisher Scientific) at -20 °C or snap frozen in liquid nitrogen and stored at -80 °C until further use.

For immunohistochemistry (IHC) or immunofluorescence (IF), mice were perfused with 40 ml sterile PBS followed by 20 ml of 4 % paraformaldehyde (PFA) solution. Brains were extracted and post-fixed in 4 % PFA for 4 h (IF) or 24 h (IHC) before being transferred to 30 % sucrose solution. After two days, IF-samples were slowly frozen to -80 °C in Tissue-Tek® O.C.T.™ Compound (Sakura Finetek Europe B.V., Alphen aan den Rijn, Netherlands) using liquid nitrogen and 2-methylbutane, whereas IHC-samples were moved to PBS + 0.1 % NaN_3_ and stored at 4 °C until further use.

### Cytokine immunoassay

Serum cytokine levels were assessed using the LEGENDplex™ Mouse Inflammation Panel (BioLegend, San Diego, CA, USA) according to the manufacturer’s instructions. Flow cytometry was performed with an LSR Fortessa instrument (BD, Franklin Lakes, NJ, USA) and data was analyzed using the LEGENDplex™ Data Analysis Software (v8.0, BioLegend).

### RNA isolation

Tissue samples were collected and stored in RNAlater solution as described above and prepared for RNA isolation as follows: Isolated brain regions were homogenized in lysis buffer using BashingBeads Lysis tubes (Zymo Research Europe, Freiburg, Germany) and isolated using AllPrep DNA/RNA Mini Kit (Qiagen, Hilden, Germany) according to the manufacturer’s instructions. Concentration and purity of isolated RNA was determined using a NanoDrop 2000 spectrophotometer (Thermo Fisher) and stored at -80 °C until further use.

### Reverse transcription qPCR

Gene expression levels of cytokines, inflammatory mediators, tight junction proteins, neurotrophins and neurotrophin receptors, synapse associated protein, and complement system were assessed in duplicates using either 30 ng isolated RNA or 600 ng for absolute quantification of viral load, respectively. Amplification was carried out with TaqMan® RNA-to-CT™ 1-Step Kit (Applied Biosystems, Foster City, CA, USA) or Power SYBR® Green RNA-to-CT™ 1-Step Kit (ThermoFisher Scientific) and LightCycler® 96 (Roche, Basel, Switzerland) as previously described [29]. Thermal-cycling parameters were set as follows: reverse transcription (48 °C, 30 min), inactivation (95 °C, 10 min) followed by 55 cycles of denaturation (95 °C, 15 s) and annealing/extension (60 °C, 1 min). In case of the SYBR® green RT-PCR, amplification was followed by a melting curve analysis. Utilized TaqMan® Gene Expression Assays (Applied Biosystems) are listed in **Table 1**. Self-designed primers were synthetized by Tib MolBiol (Berlin, Germany) and used at 100 nM final concentration (listed in **Table 2**). For relative quantification, expression of *Hprt* was chosen as reference and relative target gene mRNA levels were determined by the ratio target gene / reference gene and subsequently normalized to mean values of control group. For absolute quantification of viral load, a standard curve was established by using a reference plasmid standard with known numbers of IAV Nucleoprotein (NP) copies per sample (1.5 x 10^1^ to 1.5 x 10^9^) [30].

**Table 1:** TaqMan® Assays used for RT-qPCR analyses.

| Protein | Gene | Assay ID |
| --- | --- | --- |
| BDNF | Bdnf | Mm04230607_s1 |
| C1qa | C1qa | Mm00432142_m1 |
| C3 | C3 | Mm01232779_m1 |
| CCL2 | Ccl2 | Mm00441242_m1 |
| CD36 | Cd36 | Mm00432403_m1 |
| CD68 | Cd68 | Mm03047343_m1 |
| Claudin-5 | Cldn5 | Mm00727012_s1 |
| HPRT | Hprt | Mm01545399_m1 |
| IDO | Ido1 | Mm00492586_s1 |
| IFN-β | Ifnb1 | Mm00439552_s1 |
| IFN-γ | Ifng | Mm00801778_m1 |
| IGTP | Igtp | Mm00497611_m1 |
| IL-1β | Il1b | Mm00434228_m1 |
| IL-6 | Il6 | Mm00446190_m1 |
| iNOS | Nos2 | Mm00440485_m1 |
| NGF | Ngf | Mm00443039_m1 |
| IRGM1 | Irgm1 | Mm00492596_m1 |
| p75<sup>NTR</sup> | Ngfr | Mm01309638_m1 |
| TNF | Tnf | Mm00443258_m1 |
| TREM-2 | Trem2 | Mm04209422_m1 |
| TrkB | Ntrk2 | Mm00435422_m1 |
| VGLUT1 | Slc17a7 | Mm00812886_m1 |
| ZO-1 | Tjp1 | Mm00493699_m1 |

**Table 2:**
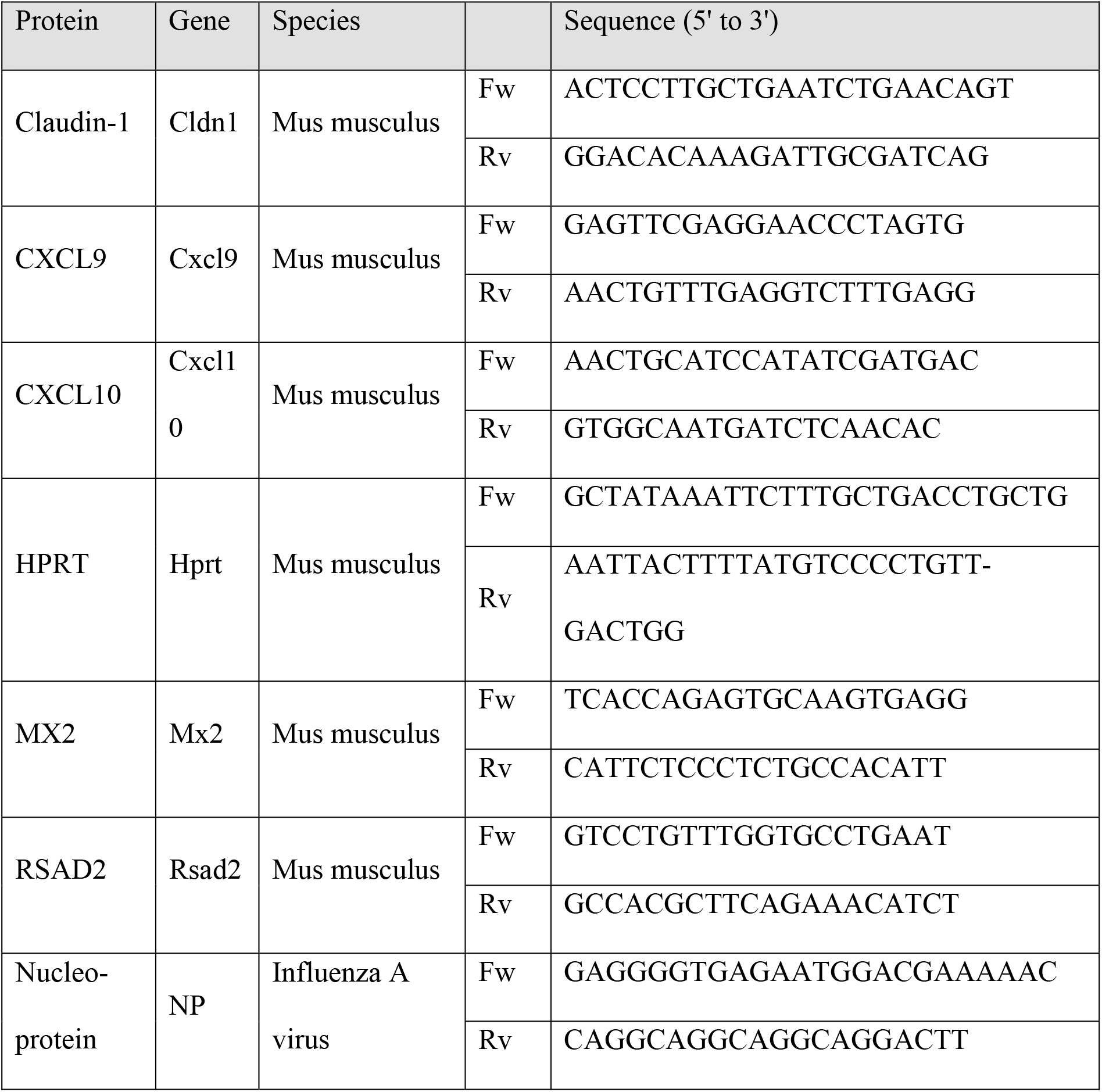
Self-designed primer sequences used for RT-qPCR analyses.

### Histopathology and immunohistochemistry

For histopathology, brains collected from naive and infected mice were first dehydrated in Zinc fixative (BD) followed incubation in ethanol at concentrations increasing from 70 % to 100 %. Then, brains were embedded in paraffin and sagittal sections of 3 µm were mounted on object slides before de-paraffinization in Xylol and alcohol at concentrations decreasing from 100 % to 70 %. For Hematoxylin and eosin staining, slides were first stained in Mayer’s hemalum solution (Sigma-Aldrich, St. Louis, MO, USA) and rinsed with tap water and 0.1 % HCl before applying eosin Y solution (Sigma-Aldrich) and rinsing in distilled water. Staining with antibodies against CD11b (dilution 1:100, #HS-384 117, Synaptic Systems, Göttingen, Germany), IBA1 (dilution 1:500, #HS-234 017, Synaptic Systems), MAP2 (2 µg/µl, #188 011, Synaptic Systems), and NeuN (dilution 1:2000, #266 008, Synaptic Systems) was performed at room temperature for 1 h after antigen retrieval using 10 mM citrate + Tween (pH 6.0) and blocking of endogenous peroxidase activity. Subsequently, slides were incubated with matching, biotinylated secondary antibodies and developed using VECTASTAIN® ABC-HRP Kit (#PK-4000, Vector Laboratories, Burlingame, CA, USA). Upon incubation with DAB substrate, samples were counterstained with Hematoxylin and rinsed with tap water, then dehydrated in alcohol at increasing concentrations, Propanol, and Xylol before mounting. Finally, sections were imaged and analyzed using an Olympus VS120 virtual-slide-microscope (Olympus Life Science, Waltham, MA, USA) equipped with an Olympus VC50 camera and a 40x objective, and Olympus VS-ASW imaging software (version 2.9.2 Build 17565, Olympus Life Science).

To obtain samples for immunohistochemistry, sagittal sections of 20 µm from frozen brain tissue were transferred to SuperFrost Plus™ (Thermo Scientific) slides. Antigen retrieval (10 mM Citrate buffer, pH 6.0, 0.1 % Tween-20) was performed at 96 °C for 30 min. Blocking and permeabilization was performed in PBS + 0.3 % (v/v) Triton X-100 with 5 % normal-goat-serum and unconjugated F(ab′)2 goat anti-mouse IgG (H+L) antibody (1:500, Thermo Scientific), at 4 °C for 2 h. Subsequently, sections were incubated with the following primary antibodies at 4 °C overnight: anti-IBA1 (dilution 1:200, #HS-234 004, Synaptic Systems) and anti-TMEM119 (dilution 1:200, #ab209064, abcam, Cambridge, UK). Next, slides were washed and subsequently incubated with matching secondary antibodies (dilution 1:1000) at room temperature in the dark for 1 h before mounting on a glass slide using ProLong™ Gold antifade mountant with DAPI (Thermo Fisher). To image microglia cells in the Cortex and Hippocampus, z-stacks were generated with a 20x objective at a z-step of 1 µm using a SP8 laser scanning confocal microscope (Leica Biosystems, Nußloch, Germany). Composite images were generated using ImageJ with Fiji distribution [31].

### Flow cytometric analysis

For flow cytometric analysis of cell phenotypes, freshly isolated cells were first incubated with Zombie NIR™ fixable dye (BioLegend, CA, USA) for live/dead discrimination and with anti-FcγIII/II receptor antibody (clone 93) to prevent unspecific binding of antibodies. Cells were further stained with the following fluorochrome-conjugated antibodies against cell surface markers in FACS buffer (PBS containing 2 % fetal bovine serum and 0.1 % sodium azide): eFluor™ 450-CD45 (clone 30-F11) and FITC-MHC Class I (clone 28-14-8), APC-CD11b (clone M1/70) (purchased from eBioscience™, San Diego, CA, USA); Brilliant Violet™ 421-CD86 (clone GL-1), Brilliant Violet™ 510-F4/80 (clone BM8), Brilliant Violet™ 605-F4/80 (clone BM8), Brilliant Violet™ 605-CD11b (clone M1/70), PerCP/Cy5.5-CD80 (clone 16-10A1), PE/Dazzle594™-MHC Class II (I-A/I-E) (clone M5/114.15.2), PE/Cy7-CX3CR1 (clone SA011F11) (purchased from BioLegend). Cells were acquired using a SP6800 Spectral Cell Analyzer (Sony Biotechnology, San Jose, CA, USA) and obtained data were analyzed using FlowJo software (version 10.5.3, FlowJo LLC, OR, USA). Fluorescence Minus One (FMO) controls were used to determine the level of autofluorescence.

### Preparation of synaptosomes from frozen brain samples

Synaptosomes were obtained from Cortex and Hippocampal formation according to protocols published elsewhere [32] but with slight modifications: After first homogenization steps, the crude membrane pellet P2 was fractioned on a discontinuous sucrose gradient with layers of 5 mM Tris/HCl pH 8.1 containing 0.32 M, 1.0 M or 1.2 M sucrose at 80,000 x g, 4 °C for 2 h. Subsequently, synaptic material was collected from the 1.0/1.2 M interphase and washed with SET buffer (320 mM sucrose, 1 mM EDTA, 5 mM Tris, pH 7.4) at 100,000 x g, 4 °C 1 h. Finally, the pelleted synaptosomes were resuspended in SET buffer containing 5 % DMSO, aliquoted and slowly frozen to - 80 °C using an isopropanol freezing container and stored until further use [33].

### Flow Synaptometry

Aliquots of frozen synaptosomes were thawed in a water bath at 37 °C and centrifuged for 10 min at 14,000 x *g* to remove sucrose-containing buffer. Supernatant was removed gently and pellets were resuspended in fixation buffer (FoxP3 Transcription Factor Staining Buffer Set, #00-5523-00, eBioscience™) and incubated on ice for 45 min. Subsequently, samples were centrifuged again for 10 min at 14,000 x *g*, resuspended in permeabilization puffer (FoxP3 Transcription Factor Staining Buffer Set, eBioscience™) reconstituted with 10 % Normal Goat Serum (NGS, ThermoFisher) and stained with primary antibodies against Gephyrin (#ab136343, abcam), GluR1 (#ABN241, Sigma-Aldrich), Homer1 (#MAB6889, R&D Systems, Minneapolis, MN, USA), Synaptophysin (#101 004, Synaptic Systems), and VGLUT1 (#135 303, Synaptic Systems). Following incubation, samples were washed and resuspended in Permeabilization buffer + 10 % NGS and stained with matching secondary antibodies: goat anti-mouse AlexaFluor® 405 (#A31553, ThermoFisher), goat anti-rabbit AlexaFluor® 488 (#ab150081, abcam), goat anti-chicken AlexaFluor® 546 (#A-11040, abcam), and goat anti-guinea pig AlexaFluor® 647 (#A21450, ThermoFisher). Finally, samples were washed once more and resuspended in SET buffer before adding the styryl dye FM4-64 (#T13320, ThermoFisher) to a final concentration of 0.2 µg/ml [33]. Samples were acquired using the Attune NxT Flow Cytometer (ThermoFisher) equipped with 405, 488, 561, and 633nm lasers. Voltages for forward-scatter light (FSC), side-scatter light (SSC) and fluorescence detection channels were set as follows: FSC 400 V, SSC 500 V, VL1 400 V, BL1 400 V, BL3 380 V, YL1 400 V, RL1 440 V. For optimal acquisition of synaptosomes, the FSC-triggered detection was replaced by a fluorescence-triggered detection with FM4-64 in the BL3 channel (threshold set to 0.3 x 10^3^ to select only FM4-64-positive events). Furthermore, the event rate was kept below 300 events/second by utilizing the slowest flow rate in combination with an adequate dilution of the sample prior to measurement to reduce coincident particle detection. A size range from 300 - 1000 nm was applied to detected events in the FSC channel using red-fluorescent silica beads with a diameter of 300 nm (#DNG-L020, Creative Diagnostics, Shirley, NY, USA) and 1000 nm (#DNG-L026, Creative Diagnostics) [33]. Obtained data were analyzed using FlowJo software (version 10.5.3, FlowJo LLC, Ashland, OR, USA). Fluorescence Minus One (FMO) controls were used to determine the level of autofluorescence.

### Western blot analysis of synaptic proteins

For Western blot analysis, aliquots of isolated synaptosomes were thawed in a water bath at 37 °C and centrifuged for 10 min at 14,000 x *g* to remove sucrose containing buffer. Pellets were lysed for 30 min at 37 °C in RIPA lysis buffer containing protease inhibitors, 50 mM Tris/HCl (pH 7.4); 150 mM NaCl; 1 % IGEPAL CA-630; 0.25 % Na-deoxycholate; 1 mM NaF. Subsequently, samples were centrifuged for 30 min at 100,000 x *g* and only supernatants were used for further separation on a 12.5 % SDS-PAGE with loading buffer (50 mM Tris/HCl pH 6.8, 100 mM dithiothreitol, 2 % SDS, 1.5 mM bromophenol blue, 1 M glycerol). Next, proteins were transferred to nitrocellulose membranes and incubated with antibodies against VGLUT1 (dilution 1:1000, Synaptic Systems) at 4 °C overnight. For the loading control, membranes were incubated with a 1:1000 dilution of anti-β-Actin antibody (#4970, Cell Signaling Technology, Cambridge, UK). Following incubation, membranes were incubated with matching HRP-conjugated secondary antibodies at room temperature for 2 h before revealing bound antibodies using enhanced chemiluminescence assay. Densitometric analysis of blots was performed using ImageJ with Fiji distribution [31].

### Electron microscopy of synaptosomes

Electron microscopic analysis of isolated synaptosomes was performed according to Breukel et al. [34] but with minor modifications: Synaptosomes were first fixed in 0.1 M cacodylate buffer (pH 7.4) with 2.5 % paraformaldehyde and 2.5 % glutaraldehyde in the refrigerator overnight. Then, the suspension was post-fixed in 0.1 M cacodylate buffer containing 1 % osmium tetroxide for 30 minutes and rinsed with distilled water subsequently. This was followed by a stepwise dehydration in a graded series of ethanol (50 % - 100 %) for 5 minutes each. Finally, the suspension was embedded in Durcupan ACM (Honeywell Fluka™, Morristown, NJ, USA) by dropping it carefully into the tubes. The resin polymerized at 70 °C for 3 days. For sectioning, the tubes were cut off and the complete block of Durcupan with the pellet of synaptosomes at the tip was directly placed in an ultramicrotome (Ultracut E, Reichert-Jung, Wetzlar, Germany). Ultrathin sections of 50-70 nm were collected on Formvar-coated slot grids of copper and examined with a LEO 912 transmission electron microscope (Carl Zeiss, Oberkochen, Germany) and imaged with a MegaScan™ 2K CCD camera (Gatan In-+c., CA, USA) using the DigitalMicrograph® software (version 2.5).

### Statistical analysis

Relative body weight was compared by multiple *t*-test with Holm-Sidac *post-hoc* correction. Data from flow cytometry and RT-qPCR were compared by Student’s *t*-test with Welch’s correction and Western blot analysis by one-way ANOVA with Holm-Sidak *post-hoc* correction using GraphPad Prism 7 (GraphPad Software, CA, USA) and R (version 4.0.3) [35] with “lattice” package [36]. Data shown is representative of three independent experiments. In all cases, results are presented as arithmetic mean and were considered significant, with *p* < .05.

## 7. Results

### Infection with influenza A virus (H1N1) alters brain homeostasis of mice and temporarily affects blood-brain barrier function

To explore the effects of virus-induced peripheral inflammation on microglia activation and neuronal alterations, we infected mice with a low dose of the mouse-adapted influenza A virus (IAV)/PR8/H1N1 and monitored their body weight and cytokine levels in serum and lung throughout the infection (**Fig. 1a, b**). After 4 to 5 days post infection (dpi) with 0.04 of the median lethal dose (MLD)_50_ of IAV, the body weight of infected mice started to decline and this trend became significant after 7 dpi (naive 100.8 ± 0.7 g *vs.* IAV-infected 89.8 ± 1.9 g, *p* < .05) while reaching its peak at 8 dpi (naive 102.2 ± 0.6 g *vs.* IAV-infected 86.7 ± 2.5 g, *p* < .001) (**Fig. 1b**). Beyond this point, mice started to recover and reached their initial weight by 14 dpi (naive 107.9 ± 0.7 g *vs.* IAV-infected 102.9 ± 1.9 g, *p* < .81). These observations corresponded well to the levels of cytokines and chemokines we detected in these mice: In the lungs, cytokines increased strongly after 4 dpi and remained elevated until 9 dpi (**Suppl. Fig. 1**). Although not reaching equally high concentrations, we also found serum levels of IFN-γ (156.0 ± 34.4 pg/ml, 6 dpi), IL-6 (37.7 ± 11.5 pg/ml, 6 dpi), and CCL2 (21.0 ± 7.3 pg/ml, 9 dpi) to be higher in IAV-infected mice, whereas GM-CSF and TNF remained unaltered (**Fig. 1c-j**). These observations match those from previous reports of our group, where the viral burden in the lungs of infected animals shows a strong decrease between 7 and 9 dpi and a complete clearance by 14 dpi [30].

**Fig. 1:**
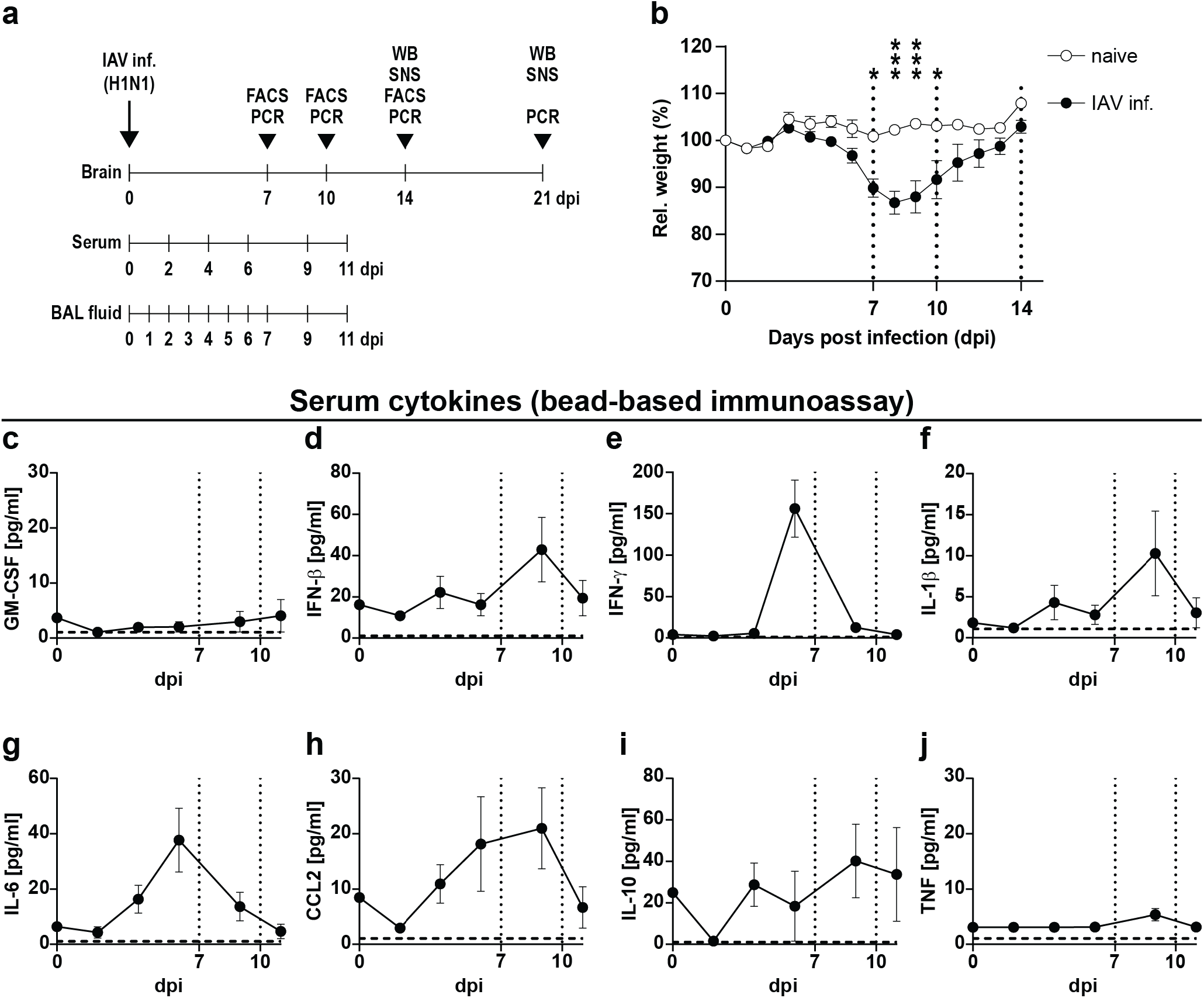
Body weight and serum cytokine levels during the course of IAV PR8/A/34(H1N1) infection. (a) Shows the experimental model used in this study. Mice were infected i.n. with a sub-lethal dose of influenza A/PR8/A/34(H1N1) and sampled between day 7 and 21 post-infection (dpi) for PCR, flow cytometry (FACS), Western blot (WB) or synaptosome (SNS) analysis. (b) Relative body weight of naive (white boxes) *vs*. infected mice (black boxes) over the course of IAV infection. Dashed vertical lines indicate the time points of experiments in line with the experimental model depicted in (a). (c-j) Serum cytokine levels in infected mice from 0 to 11 dpi. Dashed vertical lines indicate the time points of experiments in line with the experimental model and dashed horizontal lines indicate the detection limit of each cytokine, respectively. Data are shown as mean ± SEM and groups in (b) were compared by multiple Student’s *t*-tests with Holm-Sidak *post-hoc* correction. Significant differences are indicated by * (*p* < .05, *** *p* < .001).

In recent years, several studies have shown that systemic infection with pathogens or the application of pathogen-associated substances alone, such as endotoxin or poly(I:C), can induce sickness behavior and lead to disturbances in brain homeostasis [8, 12, 37, 38]. Given the high abundance of peripheral cytokines, we wondered whether this effect could also be seen in IAV-infected mice. Based on initial results, the time points of 7 dpi (first day of significantly different body weight), 10 dpi (last day of significantly different body weight), 14 dpi (restoration of initial body weight), and 21 dpi (to evaluate potential long-term effects of IAV infection) were selected for further analyses. Consequently, to evaluate the expression of cytokines and chemokines in the brains of naive and infected mice, brains were collected at the above-mentioned time points and dissected into Cortex (CTX) and Hippocampal formation (HPF). Reverse transcription qPCR (RT-qPCR) analysis revealed that upon acute IAV infection, the expression of *Il1b* (IL-1β), *Il6* (IL-6), *Tnf* (TNF), and *Ccl2* (CCL2) remained unchanged (**Fig. 2a-d**). Although previous studies have demonstrated IAV/PR8/34 to be non-neurotropic [12, 13], we evaluated the viral load in the Cortex, Hippocampus, and Olfactory bulb at 7 and 10 dpi to exclude direct viral effects in the CNS of infected mice. In line with published data, we did not detect viral copies in these brain regions (**Fig. 2e**), however, despite the absence of IAV in the CNS, mRNA levels of *Ifnb1* (IFN-β) were elevated in the Cortex at 10 dpi (*p* < .055) and significantly increased by 14 dpi. A similar trend was also observed for *Ifng* (IFN-γ) **(Fig. 2f, g**). As the induction of interferons often results in a plethora of immune responses ranging from the establishment of an anti-viral state in neighboring cells to modulation of the adaptive immunity, we sought to investigate for an altered regulation of interferon-stimulated genes (ISGs). Indeed, we found the expression of the inducible GTPase *Irgm1* (IRGM1) to be significantly increased in the Cortex 10 dpi, whereas expression of other ISGs such as *Igtp* (IGTP), *Mx2* (MX2) or *Rsad2* (RSAD2) remained unaffected (**Fig. 2h-k**). In addition to anti-viral state induction, type I and type II interferons have been recently identified as key modulators of immune cell entry to the CNS during physiological but also pathological conditions by affecting the leukocyte trafficking via the blood-brain barrier (BBB) or the blood-cerebrospinal fluid barrier (BCSFB) located in the choroid plexus [39]. Interestingly, our data revealed that the gene expression of barrier-associated tight-junction proteins *Cldn5* (Claudin-5) and *Tjp1* (ZO-1) [40] was significantly altered in the Cortex and Hippocampus upon IAV infection (**Fig. 3a-c**). However, the expression level of the chemokines CXCL9 and CXCL10, known attractants of leukocytes to the CNS [41, 42], showed no significant differences (**Fig. 3d, e**). In summary, peripheral infection with the influenza A virus led to increased gene expression of interferons in the brain combined with a reduction of tight junction proteins, suggesting an impaired functionality of the BBB and BCSFB.

**Fig. 2:**
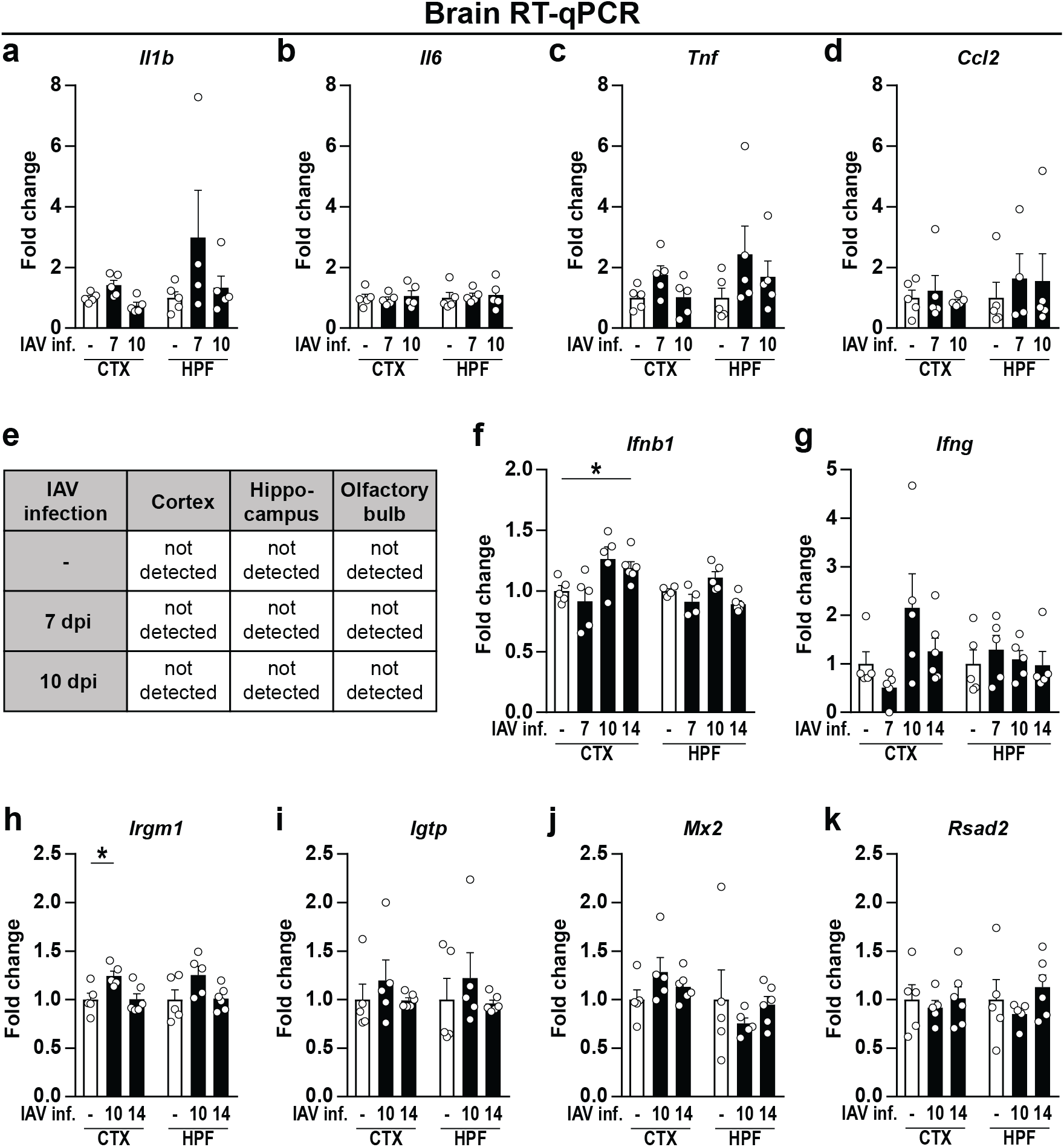
Expression level of cytokines, chemokines, and interferon-stimulated genes in brains of naive and IAV-infected mice. Brains of perfused animals were dissected into Cortex (CTX), Hippocampal formation (HPF), and Olfactory bulb (OB) according to the Allen Mouse Brain Atlas [28] and used for RNA isolation as described above. Gene expression in naive (white bars) and infected animals (black bars) is shown for (a-d) cytokines and chemokines during the acute and late phase of IAV infection (7-10 dpi); (e) IAV load in the brain during the acute and late phase of IAV infection (7-10 dpi); (f, g) type I and II interferons during IAV infection (7-14 dpi); (h-k) induction of interferon-stimulated genes during the late phase of IAV infection (10-14 dpi). Relative gene expression was examined by RT-qPCR as described above and expression of target genes was normalized to the expression level of *Hprt*. Subsequently, relative expression was normalized to the means of naive animals. Data are shown as mean ± SEM and groups were compared via Student’s *t*-test with Welch’s correction. Significant differences are indicated by * (*p* < .05).

**Fig. 3:**
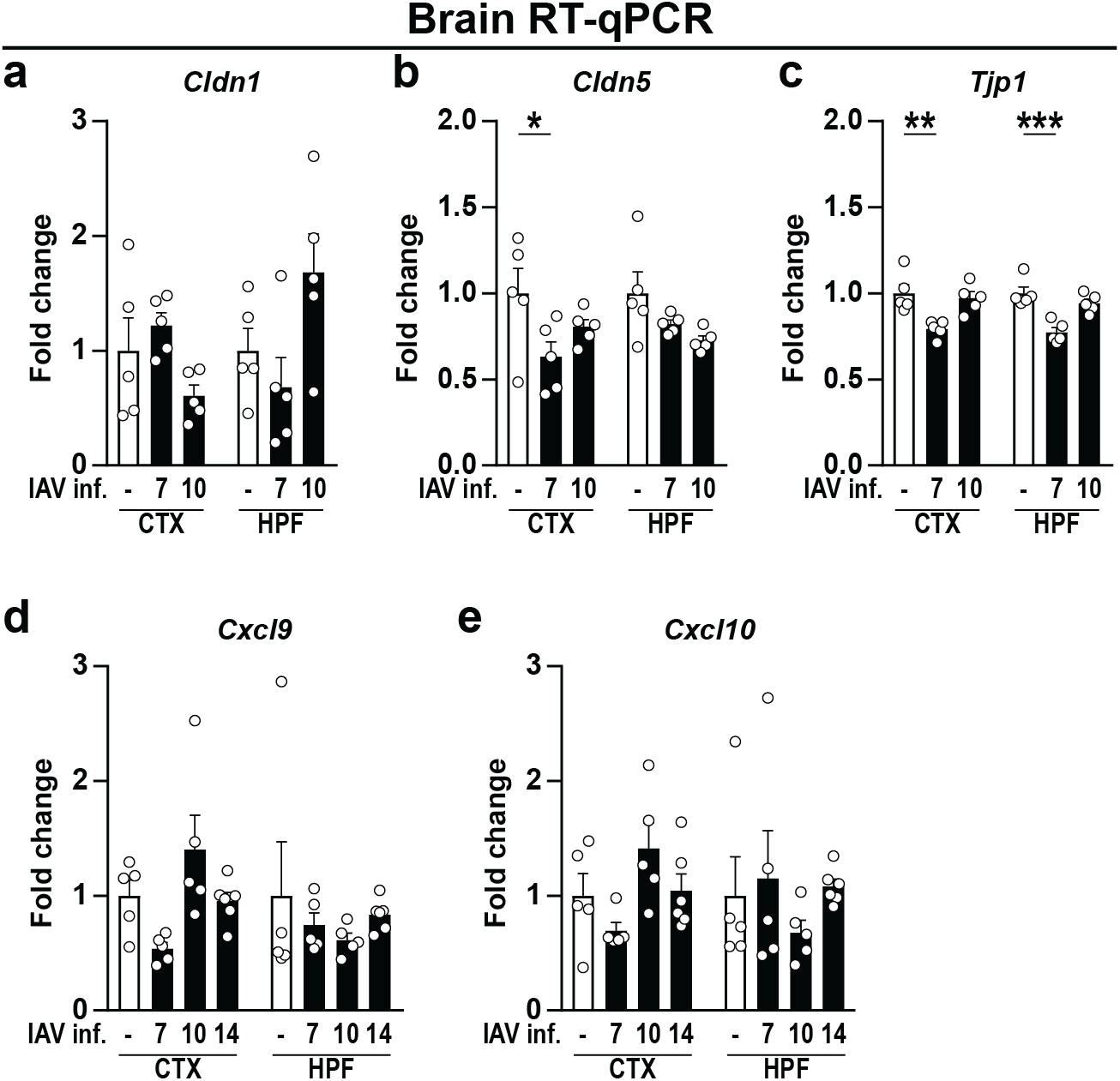
Gene expression level of blood-brain barrier-associated proteins upon infection with IAV. Brains were dissected into Cortex (CTX) and Hippocampal formation (HPF) as described above. (a-c) The gene expression levels of the tight junction proteins Claudin-1 (*Cldn1*), Claudin-5 (*Cldn5*), and ZO-1 (*Tjp1*) were examined in naive (white bars) and infected animals (black bars) during the acute and late phase of IAV infection (7-10 dpi). (d, e) Expression levels of chemokines *Cxcl9* and *Cxcl10* during IAV infection (7-14 dpi). Relative gene expression was determined by RT-qPCR as described above and expression of target genes was normalized to the expression level of *Hprt*. Subsequently, relative expression was normalized to the means of naive animals. Data are shown as mean ± SEM and groups were compared via Student’s *t*-test with Welch’s correction. Significant differences are indicated by * (*p* < .05, ** *p* < .01, *** *p* < .001).

### IAV infection results in activation of brain-resident microglia cells

Since the increased expression of cytokines can be an indicator of an induced immune response, we sought to determine whether IAV infection resulted in the emergence of microglia cell activation and neuroinflammation. Under homeostatic conditions, the majority of the brain’s immune cell population is represented by microglia cells that perform a variety of tasks to support neuronal functions and are also known producers of type I interferons [43, 44]. During steady state, microglia are characterized by their highly ramified morphology with numerous thin processes continuously surveilling the brain parenchyma. Upon encountering pathogen- or damage-associated molecular patterns, these cells become activated, retract their extensions and become highly mobile while migrating towards the sites of inflammation [15].

Thus, sagittal paraffin sections of brains from naive and IAV-infected mice were stained for ionized calcium binding adapter molecule 1 (IBA1) and the complement receptor 3 (CD11b). Compared to naive mice, IBA1-positive cells in Cortices and Hippocampi of IAV-infected animals did not differ in numbers or their ramified morphology (**Fig. 4a, c**). Similarly, histological examination of CD11b-positive cells also revealed no prominent differences from controls at 14 or 21 dpi (**Fig. 4b, d**). Furthermore, staining for neuronal markers MAP2 and NeuN did not indicate neuronal changes as results of IAV infection-induced neuroinflammation (**Suppl. Fig. 2**). Yet, IBA1 and CD11b expression is not exclusive to brain-resident microglia cells as these markers can also be expressed by infiltrating monocytes or border-associated macrophages. To confirm whether the cells observed during histopathological examination indeed represent microglia, we employed immunofluorescence microscopy of cryosections with staining for the microglia-specific marker Transmembrane protein 119 (TMEM119) [45]. Even though TMEM119 has been reported to be a marker with robust expression, its signal did not allow a detailed evaluation of microglia morphology as it was rather present on their cellular processes but not the soma (**Fig. 4e**). However, due to the substantial overlap of signal between IBA1 and TMEM119 we concluded that the majority of IBA1-positive cells in our samples constitute of microglia (**Fig. 4f**). Hence, these cryosections from naive and IAV-infected mice were further used to compare microglia morphology in more detail. Coherent with the previous histological observations, microglia in the Cortex and Hippocampus retained their ramified morphology with thin extensions after 10 and 14 dpi. In summary, histology and fluorescence microscopy revealed no obvious evidence of pathological changes in the brain following respiratory IAV infection.

**Fig. 4:**
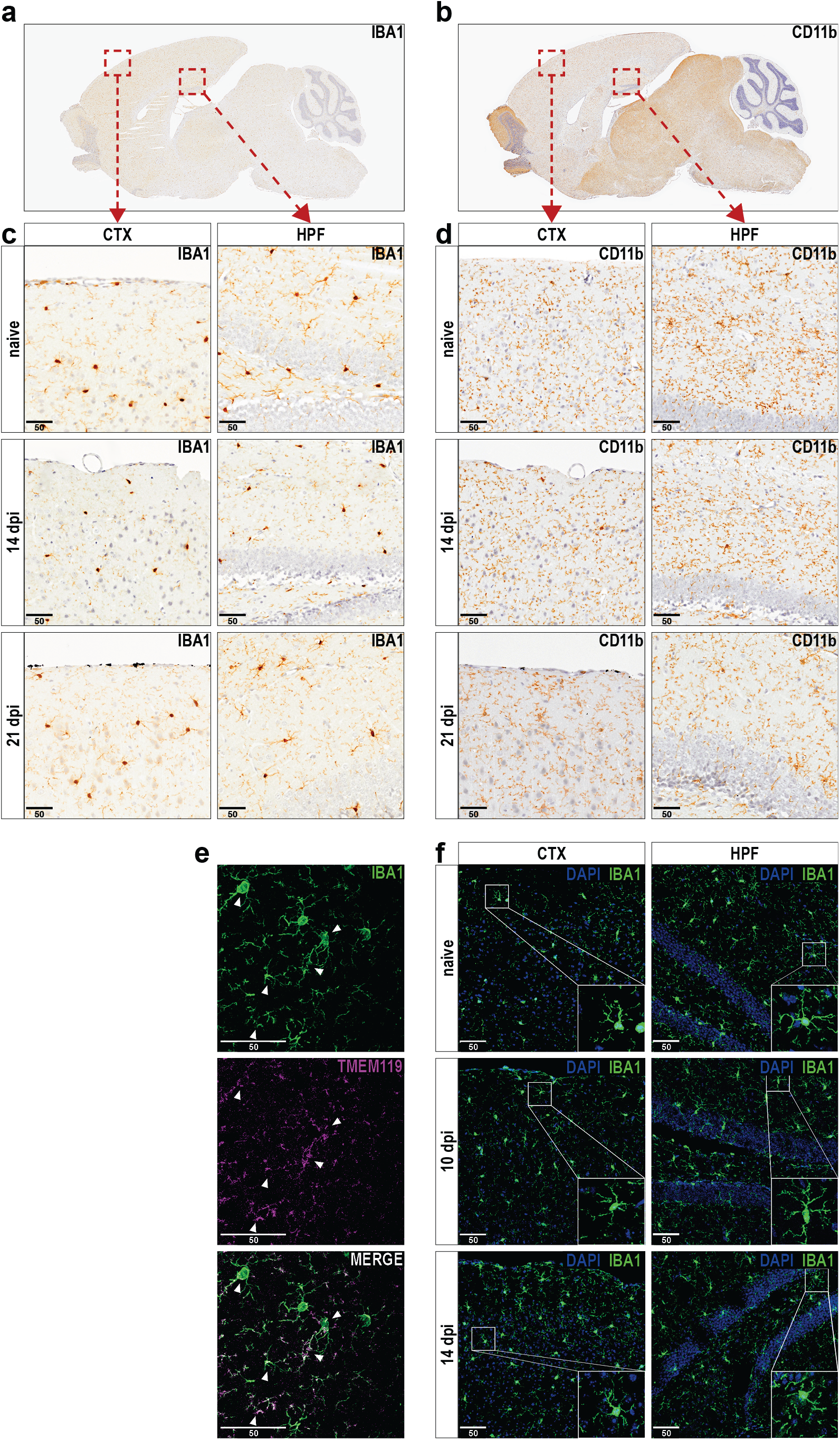
Histopathological and immunohistochemical examination of brain tissue does not reveal alterations upon infection with IAV. (a, b) Histopathological overview of representative sagittal paraffin sections from brains of naive mice stained against IBA1 or CD11b. (c, d) Panels show the Cortex (CTX, left panels) and Hippocampal formation region (HPF, right panels) from naive and infected mice (14 and 21 dpi) upon staining for IBA1 or CD11b. (e) Immunohistochemical preparation of cryosections shows representative images of microglia cells stained with antibodies for IBA1 and TMEM119 and white arrows indicate strongly overlapping signals in both channels. (f) Comparison of microglia morphology in cryosections from naive (top panels) and IAV-infected mice (middle and bottom panels) during the late phase of IAV infection (10-14 dpi). Sections were stained with antibodies against IBA1 and confocal microscopy images were generated for Cortex (CTX, left panels) and Hippocampal formation (HPF, right panels). Inserts in bottom right corners show selected microglia cells with higher detail. Scale bars = 50 µm.

Microglia were previously shown to possess a diverse phenotypic spectrum with multidimensional activation profiles that highly depend on the microenvironmental stimuli [46, 47]. Accordingly, we isolated these cells from brains of naive and IAV-infected mice to characterize their phenotype at higher resolution via multiparametric flow cytometry. After the initial removal of debris, we subjected all cells to unsupervised *t*-distributed stochastic neighbor embedding (*t*-SNE) resulting in three different clusters of cells (**Fig. 5a**) that were highly distinct in their surface expression levels of CD45, CD11b, and CX_3_CR1 and showed minor differences in the expression of major histocompatibility complex (MHC) class I and II, CD80, CD86, and F4/80. Using a subsequent manual gating approach, cells within the three different clusters were identified as brain-resident microglia cells (CD45^low^CD11b^+^) and peripheral immune cell subsets consisting of CD45^hi^CD11b^-^ and CD45^hi^CD11b^+^ cells (**Fig. 5b, c**). Unexpectedly, we detected a minor but significant increase in the frequency of CD45^high^CD11b^+^ but not CD45^hi^CD11b^-^ cells at 7 dpi in the brains of IAV-infected mice (**Fig. 5d-f**). As the infection progressed (10 dpi), not only CD45^hi^CD11b^+^ but also more CD45^hi^CD11b^-^ cells were found in the Hippocampus and Cortex (CTX *p* < .06) (**Fig. 5g-i**). Upon resolution of the peripheral infection (14 dpi), the number of CD45^hi^CD11b^+^ cells returned to levels of naive controls whereas the CD45^hi^CD11b^-^ cell population remained elevated in the Hippocampus of infected mice (**Fig. 5j-l**). However, microglia constituted by far the largest population of immune cells at all time points and thus confirmed our previous immunofluorescence results.

**Fig. 5:**
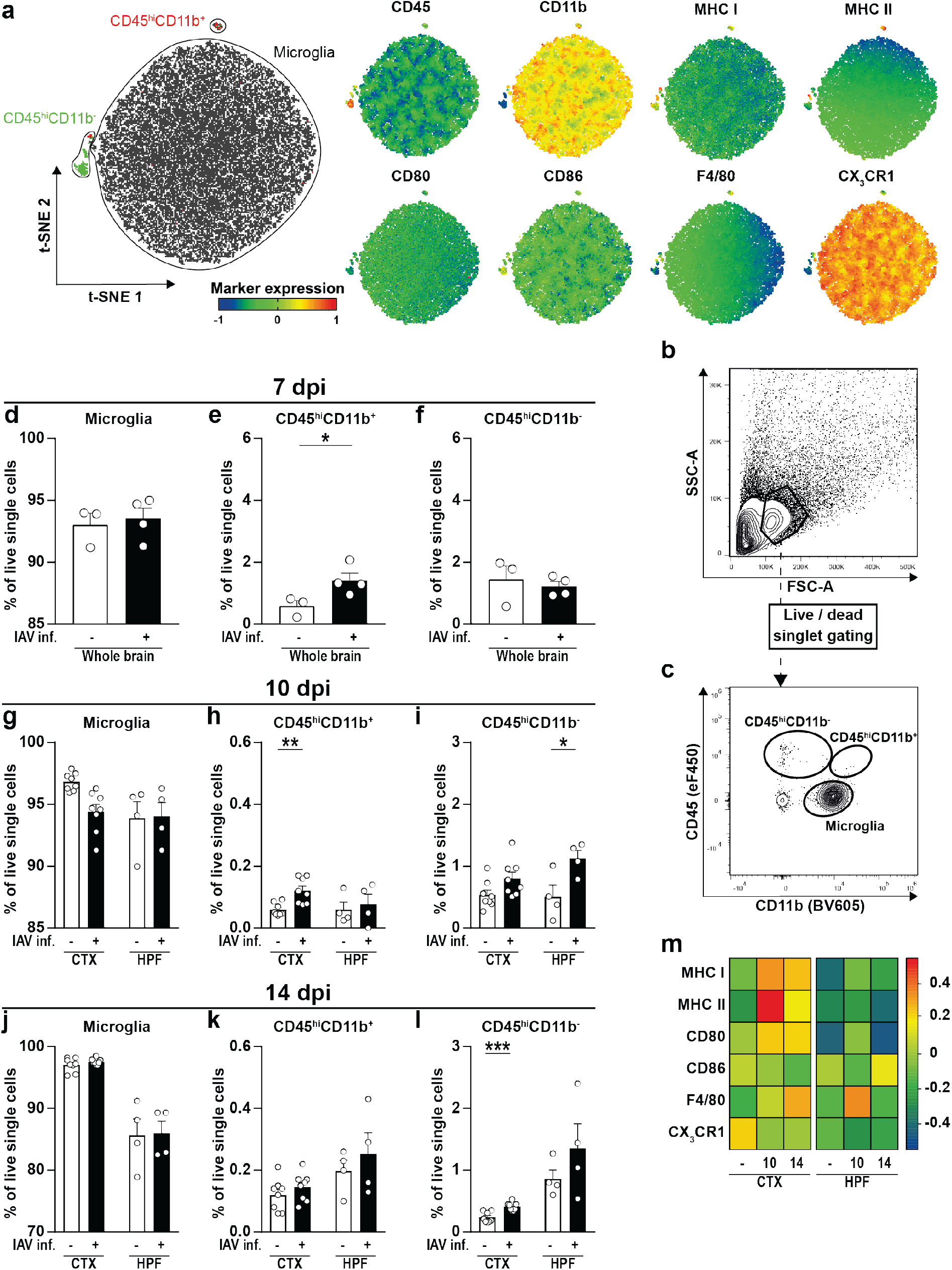
Characterization of immune cell subsets and microglia activation in the brains of IAV-infected animals. Immune cells were isolated from perfused brains of naive and infected animals as described above and subjected to flow cytometric analysis. (a) Unsupervised clustering of immune cell subsets was performed by *t*-distributed stochastic neighbor embedding (*t*-SNE) and cells within clusters were subsequently identified by manual gating. Further, differential expression of surface markers CD45, CD11b, major histocompatibility (MHC) complex class I and II, CD80, CD86, F4/80, and CX3CR1 is shown for the generated clusters. (b, c) Representative strategy for manual gating. First, cells were selected based on the forward-scatter/side-scatter plot (FSC/SSC) before exclusion of dead cells as well as doublets (not shown). Then, immune cell populations were separated by their expression of the surface markers CD45 and CD11b into brain resident CD45^low^CD11b^+^ microglia cells and recruited CD45^hi^CD11b^-^ or CD45^hi^CD11b^+^ cells. (d-f) Bar charts show the frequencies of identified immune cell populations in the brains of naive (white bars) and IAV-infected animals (black bars) at 7 dpi. (g-l) Bar charts show the frequencies of identified immune cell populations in specific brain regions (Cortex – CTX / Hippocampal formation – HPF) of naive (white bars) and IAV-infected animals (black bars) at 10 and 14 dpi. (m) Heat map plot of relative microglia surface expression of MHC I, MHC II, CD80, CD86, F4/80, and CX3CR1 in naive and infected mice at 10 and 14 dpi. Median fluorescence intensities of expressed markers were normalized to their overall mean, respectively, and data was plotted using R with “lattice” package. Data are shown as mean ± SEM and groups were compared via Student’s *t*-test with Welch’s correction. Significant differences are indicated by * (*p* < .05, *** *p* < .001).

After identification of microglia, we subsequently analyzed changes in their surface marker expression upon IAV infection. We discovered an upregulation of several immunological molecules in both Cortex and Hippocampus 10 dpi (**Fig. 5m & Suppl. Fig. 3**): Microglia of infected mice expressed higher levels of MHC I and II, CD80, and F4/80 whereas expression of the fractalkine receptor CX_3_CR1 was reduced. At 14 dpi, microglial activation was still evident in both brain regions examined with significantly increased expression of MHC I and F4/80 and decreased expression of CX_3_CR1, respectively. However, the alterations were not as pronounced as before, suggesting a return to baseline levels after resolution of peripheral IAV infection. Taken together, flow cytometric analysis supported our previous findings of an altered BBB by revealing an elevated number of recruited peripheral immune cells in the brain parenchyma. Furthermore, activation of brain-resident microglia was increased upon IAV infection and remained high throughout the course of the infection.

### Altered synaptic pruning upon IAV infection-induced microglial activation

Microglia shape neuronal connections via pruning of excessive synapses [48, 49], a process which several studies have highlighted to be a hallmark of different neurological disorders [50–52]. Of note, it has been previously demonstrated that influenza infection caused cognitive dysfunction and led to an altered neuronal architecture in mice [12]. Following the aforementioned activation of microglia, we tested whether IAV infection further reactivates microglia-mediated synaptic pruning by analyzing the expression of phagocytosis-associated receptors in the Cortex and Hippocampus (**Fig. 6**). While the gene expression of the scavenger receptor *Cd36* and lysosomal-associated protein *Cd68* was unaffected at 10 dpi, their expression was significantly increased upon sustained pro-inflammatory triggers (14 dpi) (**Fig. 6a, b**). Since aberrant synaptic pruning further requires the upregulation of complement components to tag synapses [53], we consequently examined the expression of *C1qa* and *C3* which likewise increased 14 dpi in Cortex and Hippocampus, providing the prerequisites for synapse elimination (**Fig. 6e, f**). In contrast, mRNA levels of *Trem2*, an innate immune receptor implicated in cell activation and phagocytosis [54], and the inducible nitric oxide synthase (*Nos2*) remained unaffected upon IAV infection (**Fig. 6c, d**). In conclusion, the data presented suggest the dysregulation of synaptic pruning and altered synaptic function following IAV infection.

**Fig. 6:**
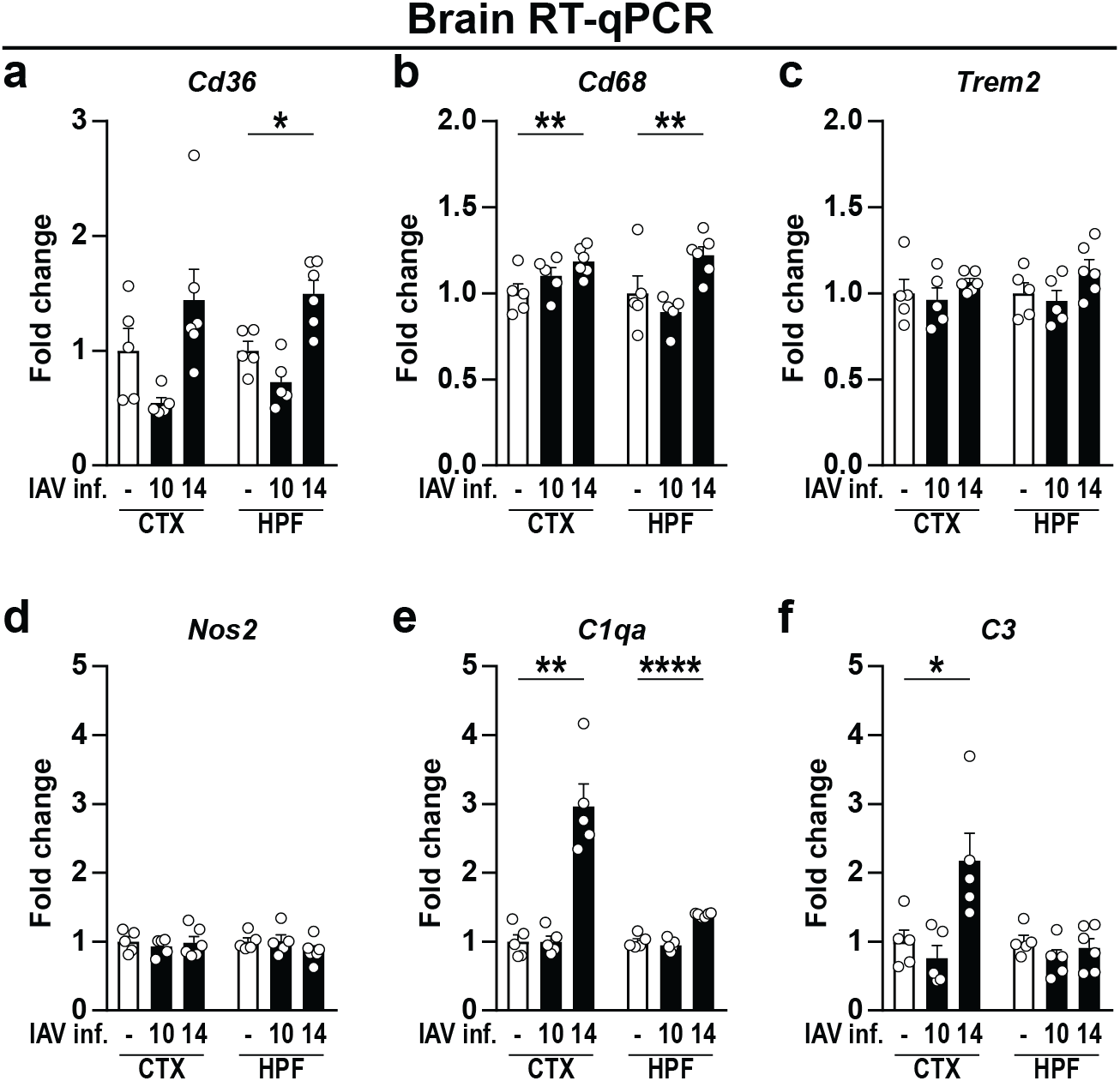
Gene expression levels of microglia activation-related genes and complement factors in brains of naive and IAV-infected mice. Gene expression in naive (white bars) and infected animals (black bars) during the late phase of IAV infection (10-14 dpi) is shown for (a-d) microglia activation-related genes and (e, f) complement factors. Relative gene expression was examined by RT-qPCR as described above and expression of target genes was normalized to the expression level of *Hprt*. Subsequently, relative expression was normalized to the means of naive animals. Data are shown as mean ± SEM and groups were compared via Student’s *t*-test with Welch’s correction. Significant differences are indicated by * (*p* < .05, ** *p* < .01, **** *p* < .0001).

### Temporally dysregulated glutamatergic synaptic transmission and neurotrophin gene expression following resolution of peripheral IAV infection

Previously, we have demonstrated that infection-induced neuroinflammation affects synaptic protein composition with detrimental outcome for glutamatergic neurotransmission [55]. To determine whether infection with IAV may affect synaptic protein composition, we analyzed the gene expression of the presynaptic glutamate transporter *Slc17a7* (VGLUT1) over the course of infection (**Fig. 7a)**. When compared to naive samples, gene expression did not differ significantly at 10 and 14 dpi, however, mRNA levels pointed towards a reduction at 14 dpi (*p* < .29). When assessing the expression levels at 21 dpi, we detected not only a return to baseline expression but significant overcompensation. Thus, we concluded that the systemic inflammation driven by peripheral IAV infection causes a functional disturbance in excitatory neurons in mice, an assumption well in line with previous reports showing altered neuronal morphology and cognitive impairment following IAV infection [12, 13]. To study synaptic changes of excitatory neurons more in-depth, we directly analyzed glutamatergic synapse composition. We therefore developed a refined approach by taking advantage of synaptosomes, *i.e.* sealed presynaptic nerve terminals often containing opposite postsynaptic elements, thus providing a well-established model to investigate synapses stripped from their surrounding tissue. Protocols to isolate synaptosomes have been existent for several decades and describe the purification of synaptic material from brain tissue using discontinuous density gradient centrifugation. Upon adaption of these protocols to our samples, we first examined the content of our isolates via electron microscopy (**Fig. 7b, c**) and detected single fragments of membranes and intact presynapses adjacent to thickened postsynapses. The imaged synaptosomes show diameters of 350-700 nm at the presynaptic side, contain zero to one mitochondrion and many small, clear synaptic vesicles. In addition, postsynaptic densities and the synaptic cleft are well noticeable, thus allowing the conclusion that our technique ensures the isolation and purification of synaptosomes from brain tissue. Secondly, we isolated proteins from our synaptosome samples and determined the protein levels of VGLUT1 by Western blot analysis before and after IAV infection (**Fig. 7d, e**). Here levels of VGLUT1 were partially diminished at 14 dpi (*p* < .09) and significantly reduced 21 dpi, thus confirming previous findings by RT-qPCR.

**Fig. 7:**
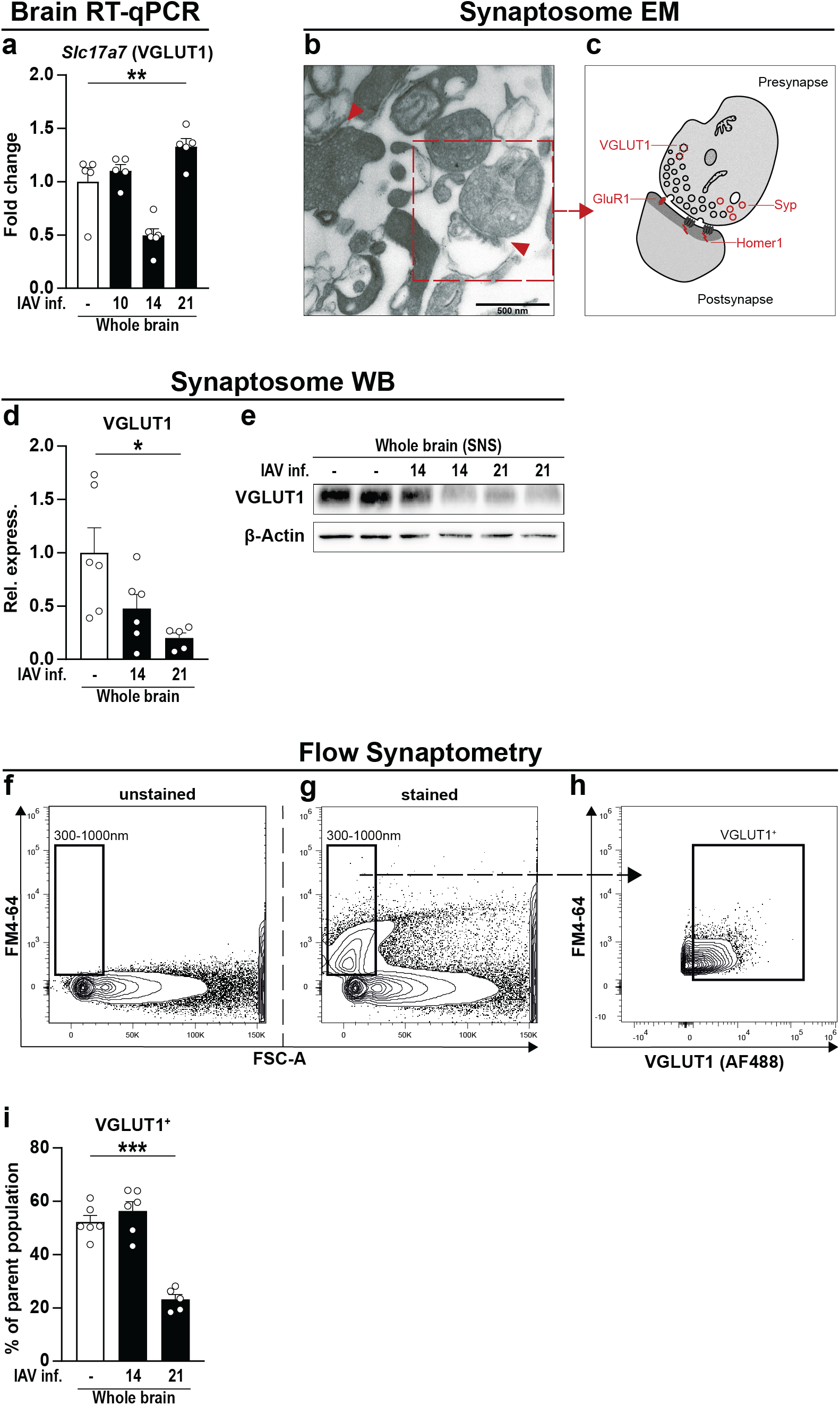
A new tool to analyze synaptic proteins via flow cytometry. (a) Gene expression VGLUT1 (*Slc17a4*) was analyzed in whole brain homogenate of naive (white bars) and IAV-infected mice at 10, 14, and 21 dpi (black bars). Relative gene expression was examined by RT-qPCR as described above and expression of the target gene was normalized to the expression level of *Hprt*. Subsequently, relative expression was normalized to the mean of naive animals. Groups were compared via Student’s *t*-test with Welch’s correction. (b) Synaptosomes were isolated from brain regions of perfused animals as described above and the synaptosomal fraction [32] was subjected to electron microscopy. Red arrows indicate intact synapses formed by a pre- and post-synaptic compartment. Scale bar in bottom left corner indicates 500 nm (c) Graphical illustration of synaptosome selected in (b). Pre-synaptic nerve endings still contain synaptic vesicles and mitochondria whereas post-synaptic domains are characterized by their region of high post-synaptic density (PSD). (d, e) Protein content of VGLUT1 from synaptosomes of whole brain homogenate was assessed via Western blot. Bar charts show the relative optical density of the proteins band from naive (white bars) and infected animals (black bars) at 14 and 21 dpi upon normalization to β-Actin expression. Values were further normalized to the mean of naive animals. Groups were compared via one-way ANOVA with Holm-Sidak *post-hoc* correction. (f-h) Isolated synaptosomes were subjected to flow cytometry and the used representative gating strategy is shown. (f) First, A gate with a size range from 300-1000 nm was established in the FSC channel using silica beads. (g) Second, separation of biological particles from residues in the buffer was facilitated using the styryl dye FM4-64 that integrates into the lipid membranes of biological organelles. Then, FSC-triggered detection was replaced by fluorescence-triggered detection with FM4-64 in the BL3 channel with a fluorescence threshold set above the noise at 0.3x10^3^ (not shown). (h) Lastly, events detected in the size range of 300-1000 nm were further gated for their expression of VGLUT1. (i) Bar chart shows the frequency of VGLUT1^+^ events in the brains of naive (white bars) and IAV-infected animals (black bars) at 14 and 21 dpi. In all cases, data are shown as mean ± SEM and significant differences are indicated by * (*p* < .05, ** *p* < .01, *** *p* < .001, **** *p* < .0001).

So far, synaptosomes have mostly been analyzed in batches [33] and only few studies are known to employ quantitative approaches [56, 57]. To investigate synapse composition at the single synapse level, we established Flow Synaptometry, a novel, flow cytometry-based approach allowing a high-throughput analyses. Reportedly, synaptosomes are rather small objects with diameters from 0.5-2 µm [58, 59]. To allow the size discrimination of detected events in a flow cytometer, we first established a size gate ranging from 0.3-1 µm by utilizing fluorescent silica beads with defined diameters (**Fig. 7f**). Typically, conventional flow cytometers display a poor FSC resolution for events of such small scale. This cannot be compensated by increasing the detector sensitivity alone as this also favors the amplification of other buffer-residing objects small enough to pass through the 0.22 µm pores of conventional sterile filters. Thus, we employed the lipophilic styryl dye FM4-64 that becomes highly fluorescent when bound to lipid bilayers, enabling us to distinguish cellular components from inorganic residues in the buffer and noise (**Fig. 7g**). This step was coupled to the replacement of the standard FSC-triggered detection by a fluorescence-triggered detection in the BL3 channel of our flow cytometer (not shown), now favoring only FM4-64-stained events while neglecting everything below the threshold. Besides increasing the flow cytometer’s sensitivity for small events, this procedure also reduces the chance of falsely detecting aggregates of multiple synaptosomes as one event [33]. Finally, we compared the frequencies of events positive for VGLUT1 (**Fig. 7h, i**) and consistently detected a significant reduction for this protein at 21 dpi, whereas no differences appeared 14 dpi. In the light of these substantial changes, we aimed to narrow down the synaptosome analysis on Cortex and Hippocampus of IAV-infected mice (**Fig. 8**). Therefore, the staining panel was expanded by specific markers that, in addition to the presynapse, verify the presence of postsynapses (**Fig. 8a, b**), thus allowing the simultaneous differentiation of intact synaptosomes from excitatory (Homer1) and inhibitory synapses (Gephyrin), respectively. When compared to naive mice, no differences in the frequency of excitatory Syp^+^Homer1^+^ or inhibitory Syp^+^Gephyrin^+^ synaptosomes were observed in animals at 14 dpi (**Fig. 8c, d**). At 21 dpi, Syp^+^Homer1^+^ synaptosomes appeared with higher frequency in Cortices of infected mice, whereas Syp^+^Gephyrin^+^ synaptosomes were found in greater percentage in the Hippocampus. Finally, the fractions of excitatory synapses positive for either the glutamate transporter VGLUT1 or the α-amino-3-hydroxy-5-methyl-4-isoxazolepropionic acid (AMPA) receptor subunit GluR1 were determined among the population of Syp^+^Homer1^+^ synaptosomes. Here results from IAV-infected animals were consistent with previous findings and did not differ from naive controls at 14 dpi, but displayed a substantial loss of Syp^+^Homer1^+^VGLUT1^+^ synaptosomes in the Cortex and Hippocampus at 21 dpi (**Fig. 8e**). In contrast, frequencies of Syp^+^Homer1^+^GluR1^+^ synaptosomes from either Cortex or Hippocampus did not appear significantly altered in the course of an IAV infection, although minor fluctuations were detectable.

**Fig. 8:**
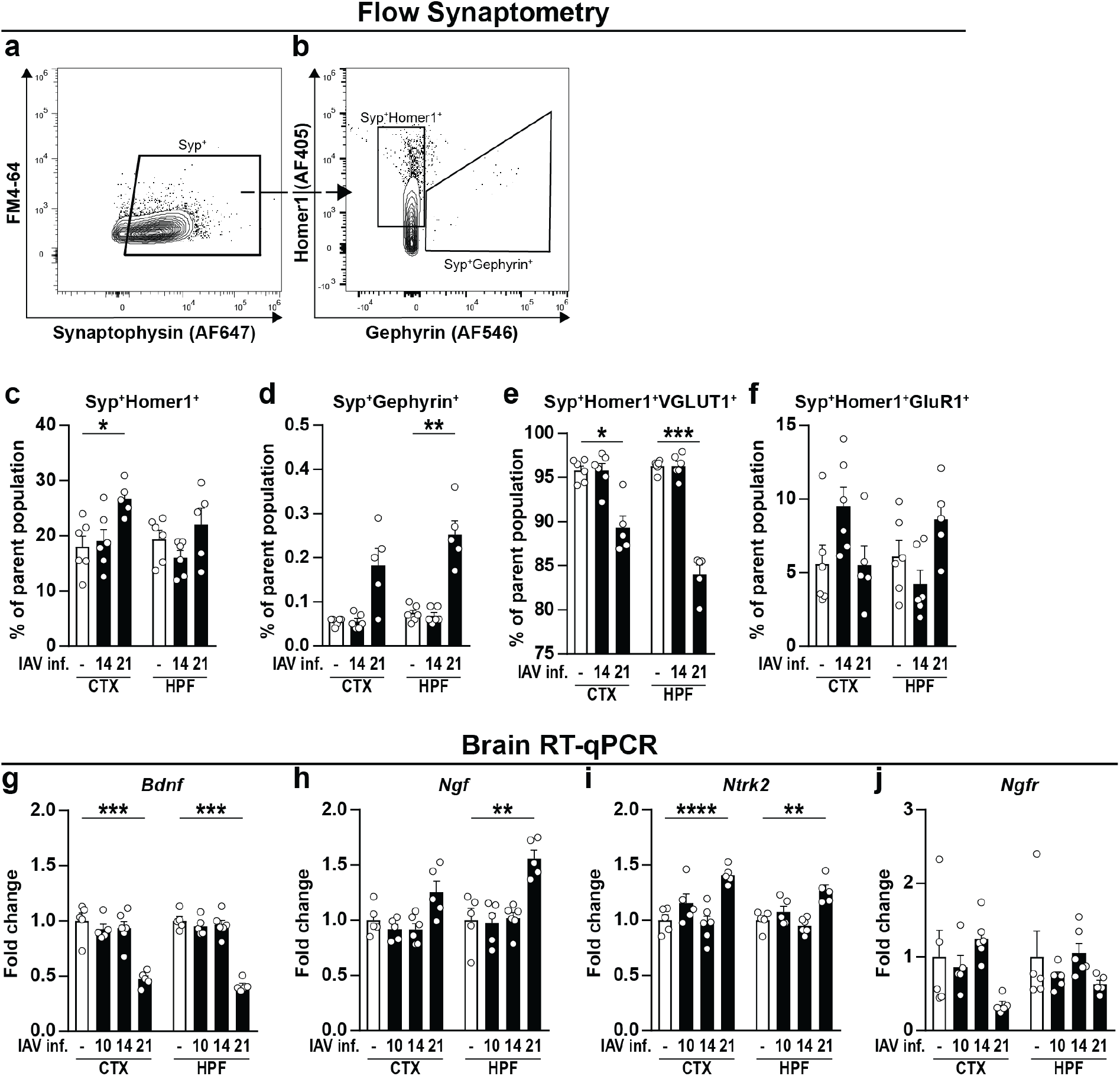
Flow cytometric synaptosome analysis and gene expression of neurotrophins and neurotrophin receptors in naive and IAV-infected mice. (a-f) Synaptosomes were isolated from Cortex (CTX) and Hippocampal formation (HPF) of naive and IAV-infected mice and subjected to further flow cytometric analysis as shown in figure 7 above. (a, b) After size gating and removal of unspecific events via fluorescence-triggered detection (not shown), Synaptophysin^+^ (Syp^+^) events were selected and subsequently gated for Homer1^+^ or Gephyrin^+^, respectively. (c, d) Bar charts show the frequencies of Syp^+^Homer1^+^ and Syp^+^Gephyrin^+^ subpopulations from naive (white bars) and IAV-infected animals (black bars) at 14 and 21 dpi. (e, f) Intact synaptosomes from excitatory synapses consisting of a pre- and postsynaptic terminal (Syp^+^Homer1^+^) were further examined for their expression of VGLUT1 and GluR1. (g-j) RNA was isolated from brains of perfused animals as described above. Relative gene expression levels of BDNF (*Bdnf*), NGF (*Ngf*), TrkB (*Ntrk2*), and p75^NTR^ (*Ngfr*) were examined by RT-qPCR on different time points post IAV infection (10-21 dpi). Expression of target genes was normalized to the expression level of *Hprt*. Subsequently, relative expression was normalized to the means of naive animals. For all graphs, data are shown as mean ± SEM and groups were compared via Student’s *t*-test with Welch’s correction. Significant differences are indicated by * (*p* < .05, ** *p* < .01, *** *p* < .001, **** *p* < .0001).

Finally, we addressed the question of whether the evident synaptic changes might, at least in part, be the result of impaired neurotrophin levels. These proteins act as important mediators of neuronal survival and differentiation in the CNS and therefore play a crucial role during the development of the brain and synaptic plasticity [60, 61]. Hence, Cortices and Hippocampi of naive and infected mice were compared by RT-qPCR at different time points post IAV infection. The levels of brain-derived neurotropic factor (*Bdnf*) and nerve growth factor (*Ngf*), both important mediators of neuronal differentiation and survival, did not vary until the late phase of IAV infection but changed significantly at 21 dpi (**Fig. 8g-j**). While expression of *Bdnf* showed a substantial reduction in Cortex and Hippocampus, *Ngf* was increased. Furthermore, the expression of the BDNF-specific neurotrophin receptor tropomyosin-receptor kinase B (TrkB / *Ntrk2*) was elevated in both brain regions 21 dpi. The pan-neurotrophin receptor p75^NTR^ (*Ngfr*), known to be differentially expressed during various neurodegenerative diseases [62, 63], remained unaltered throughout the course of IAV infection.

In summary, our data show that mRNA and protein levels of synapse components were altered upon respiratory IAV infection in mice. Furthermore, we established a novel approach to quantify the composition of synapses in brains by applying flow cytometry onto antibody-labelled synaptosomes. Thus, we highlighted that upon IAV infection the frequency of inhibitory synapses is increased while glutamatergic neurotransmission is impaired. Consistently, gene expression of neurotrophins and their receptors was also altered in the brains of infected animals 21 dpi as a possible consequence of a disturbed neurotransmission.

## 8. Discussion

Infections with influenza A virus affect all age groups and accumulate in annual epidemics. Usually, the infection is associated with fever, cough, and a sore throat but also induces symptoms of sickness behavior such as weakness or decreased interest in surroundings [64]. Moreover, cases of behavioral alterations in the form of narcolepsy [65] and the development of major depression [66] have been connected with IAV infection while other studies showed impaired hippocampal neuron morphology and cognition in the presence of neuroinflammation in mice [12, 13]. Overall, various findings suggest a link between peripheral infection and the development of neuropsychiatric disorders, however, the underlying mechanisms remained largely elusive. To address this question, we infected mice with a low dose of IAV that aimed to mimic the disease progression in humans and monitored an elevated concentration of cytokines in lungs and sera during the first days of infection. Particularly, we detected increased levels of IL-6 and IFN-γ between 6 and 9 dpi, which coincided with a prominent loss of body weight. These data are in line with our previous observations [30] and are the result of a cellular and humoral immune response facilitated by cytotoxic CD8^+^ T cells, and IFN-γ-producing NK and CD4^+^ T helper cells that leads to diminished viral burden in the lungs [7]. Recently, the induction of peripheral inflammation through administration of LPS, poly(I:C) or IAV has been shown to cause elevated serum levels of IL-6 and TNF, which led to increased expression of inflammatory cytokines in the brain [12, 22, 67, 68]. Consequently, we assessed mRNA levels of various inflammatory cytokines but were unable to detect prominent changes for *Il1b*, *Il6*, *Tnf* or *Ccl2*. Although our data and other previous studies could show that IAV PR8/A/34(H1N1) lacks the ability to infect the brain [12, 13], we found a moderate but significant increase of type I interferons (*Ifnb1*) and the IFN-γ-inducible GTPase *Irgm1* in the Cortex of infected mice at 10 dpi. Possibly, this observation may be a consequence of activated microglia in response to high serum titers of inflammatory cytokines [43, 44] infiltrating the CNS via the BBB [9, 10, 69]. However, the magnitude of *Ifnb1* expression levels remained considerably low and did not lead to a more restricted permeability of the BBB as observed in previous studies [70, 71]. In contrast, we found a diminished expression of genes associated with tight junction proteins of the BBB [40] at 7 dpi, shortly after serum levels of IFN-γ peaked in IAV-infected mice. The dysregulated state of BBB and BCSFB was further indicated by the marginally increased frequencies of peripheral immune cells we detected in Cortex and Hippocampus between 7 and 14 dpi. These results are in line with several reports demonstrating the ability of type II interferons to downregulate the expression of tight junction proteins [72, 73]. Although IFN-γ is further able to modify the expression of chemokines CXCL9 and CXCL10 by the BBB and BCSFB [74, 75], gene expression of *Cxcl9* or *Cxcl10* in the brain homogenate of IAV-infected mice revealed no difference in our studies in contrast with a previous report on poly(I:C)-induced release of CXCL10 by brain endothelia cells. In the latter study the authors associated the induction of sickness behavior in mice with BBB- and BCSFB-derived CXCL10, further leading to aggravated hippocampal synaptic plasticity via CXCR3 signaling in neurons [38]. Of note, both poly(I:C) and IAV infection result in RIG-I signaling cascade induction. However, only IAV is able to avoid immune responses by directly interfering with cellular signaling pathways or host gene expression that ultimately disturb induction of interferons and antiviral proteins [76–78]. Thus poly(I:C)-induced inflammation can only partially translate into effects observed in our IAV infection model and therefore further investigation on the transcriptome and the contribution of brain endothelia and epithelia cells to chemokine release upon IAV infection is required.

Despite the altered cytokine expression in the brain pointing towards the occurrence of neuroinflammation upon peripheral infection with IAV, histological examination did not show inflammatory foci in Cortices and Hippocampi of infected mice. In this respect, microglia number and their ramified morphology remained unaltered throughout the course of IAV infection which partially collides with data of Jurgens et *al.* [12] showing increased IBA1 staining and reduced ramification of microglia already at 7 dpi. However, those findings may just be the result of a different genetic background (BALB/c *vs.* C57BL/6JRj) and a higher IAV inoculation dose as both loss of body weight and locomotive activity of mice appeared at least 2 days earlier than in our studies, using only 0.04 MLD50 of IAV to induce the infection. Presumably, the immune system responds proportionally to the infection dose and in our experiments, animals had only a mild loss of body weight and did not succumb to the infection. However, whether the indirect activation of microglia by peripheral challenge is dose-dependent is an interesting point that awaits further elucidation. Regardless, microglia activation can appear on multidimensional scale [46, 47] and in contrast to morphologic analyses, detailed investigation via flow cytometry allowed us to detect a moderate but significant region-specific activation indicated by increased expression of MHC I and F4/80 at 10 dpi and again at 14 dpi. These observations point towards a delayed onset of neuroinflammation following a peripheral immune response. Moreover, our data suggest that IAV infection induces a rather subtle and short-term inflammation in the brain, which might remain unnoticed using common histopathological approaches. Indeed, when mice infected with IAV PR8/A/34(H1N1) were assessed for possible long-term effects at 30 dpi, no signs of neuroinflammation were apparent at all [13]. In this context, a growing body of evidence has emphasized the connection between inflammation-induced activation of microglia and the recurring pruning of synapses in neurological disorders [50–52]. In addition to the prerequisite of activated microglia, synaptic pruning is depending on components of the complement system which tag synapses for a possible elimination [53, 79]. In fact, we found increased expression of phagocytosis-related genes as well as upregulated mRNA levels of C1QA and C3 in IAV-infected animals. Interestingly, these results were apparent only at 14 dpi indicating once again temporary repercussions in the nervous system upon peripheral inflammation. In this regard, our group has previously shown the effects of neuroinflammation on synapse integrity [55, 80]. Given the reduced expression of genes associated with synapse neurotransmission we decided to test for adverse effects of IAV infection on synapse integrity. Establishing a novel flow cytometry-based approach to quantify the composition of synaptosomes derived from Cortices and Hippocampus allowed us to detect compelling effects on presynaptic glutamatergic signaling in Cortices and Hippocampi at 21 dpi while the fraction of inhibitory synapses was increased.

To date, investigation of isolated synaptosomes represents a convenient tool to study the function and physiology of synapses freed from their surrounding environment [58]. However, only few studies have utilized flow cytometric approaches to assess synaptosomes and in most cases only make use of one synaptic component [57, 81, 82]. Other reports focused on sorting of synaptosomes derived from fluorescent neurons [56] or applying mass cytometry and mass-coupled antibodies [83] but reporter mouse models or mass cytometers are not always featured equipment of a neuroscientific laboratory. Thus, our profound technique represents an easy-to-use alternative to previous approaches as we demonstrate that simple multiplexing is feasible using a conventional isolation procedure and standard flow cytometer. To our knowledge, our study is the first to extend previous attempts of Flow Synaptometry by simultaneously quantifying the relative abundance pre- and postsynaptic components in synaptosomes from different types of neurons from different brain regions. Moreover, choosing Flow Synaptometry over conventional Western blot enables to focus specifically on intact synapses, resulting in a high sensitivity while requiring only small amounts of sample material. In addition, high-throughput acquisition allows the characterization of a large number of samples within the same session and thus facilitates a high robustness of data compared to Western blot analysis.

So far, little is known about the direct consequences of peripheral infections or inflammation on glutamatergic signaling, but models using the viral mimic poly(I:C) discovered changed extracellular glutamate levels and synaptic transmission [84]. Furthermore, certain analogies can be found with respect to pathophysiological inflammation-induced depression. Here, low levels of the neurotransmitter serotonin are assumed to play a major role and rely mainly on tryptophan availability which is in turn limited by the activity of indoleamine 2,3-dioxygenase (IDO) as a part of the kynurenine pathway [85, 86]. IDO is inducible via IFN-γ, IL-1β, and IL-6 [87–89], or upon IAV infection [90]. Degradation of tryptophan occurs through IDO in peripheral organs such as liver, intestine and spleen [91] but also in brain-resident microglia, astrocytes or recruited monocytes [64, 92]. Altered levels of serotonin have been associated with a modulation of glutamatergic and GABAergic neurotransmission by altering the release of glutamate and GABA at the presynaptic side while further suppressing long-term potentiation (LTP) via inhibition of N-methyl-D-aspartate receptor (NMDA) glutamate receptor activation [93]. Although we did not detect an upregulation of IDO mRNA levels in the brains of IAV-infected mice (not shown), studies on models of LPS-induced depressive-like behavior pointed towards a considerable role of the kynurenine pathway metabolites rather than altered brain levels of serotonin [94]. Blood-derived metabolites of the kynurenine pathway can enter the brain via large neutral amino acid transporter and cause excitotoxic effects [95]. For example, levels of quinolinic acid are elevated upon peripheral inflammation [64] and functions as NMDA agonist and is found responsible for LPS-induced depression [96]. Thus, the observed change in glutamatergic neurotransmission in our study might be a consequence of dysregulated serotonin levels or the underlying tryptophan metabolism. Next to adverse effects on glutamatergic neurotransmission, IAV infection led to an altered expression of neurotrophins and their receptors. Previously, reduced expression levels of BDNF, GDNF have been discovered in brains of IAV-infected mice [12, 97], and models of poly(I:C) and LPS administration demonstrated reduced expression of BDNF, TrkB, and NGF in Hippocampus and Cortex of mice [22, 98]. On the one hand, reduced gene expression of BDNF has been associated with the development of depressive symptoms [99] and has been shown to modulate glutamatergic synaptic transmission by affecting presynaptic Ca^2+^ levels and glutamate release as well as phosphorylation of postsynaptic NMDA receptors [100]. On the other hand, studies have shown that glutamate receptor activity has a reciprocal effect on neurotrophin production, hence indicating a strongly intertwined relationship [22]. Notably, inflammation-mediated dysbalance of glutamatergic signaling has been described as a central mechanism in several neurologic disorders such as schizophrenia [101], autism spectrum disorders [102], obsessive-compulsive disorder [103] or mood disorders, and further as a comorbidity in atherosclerosis or rheumatoid arthritis [104]. Overall, this study shows that peripheral IAV infection is followed by temporal effects in the brain, however, without direct manifestation in neurodegenerative processes, as shown by our histopathological examination of neuronal markers. Nevertheless, our findings contribute to the understanding how peripheral inflammation affects CNS integrity. Of note, re-infection of humans with influenza is likely to occur in intervals of 10 to 20 years [1] and may culminate in neurological implications [105]. In this respect, a growing body of evidence supports the connection between peripheral infections and epigenetic changes in microglia which in turn render these cells to react more pro-inflammatory [106]. As a consequence, synergistic effects of multiple infections acquired throughout life could lead to a previously unappreciated contribution in the development of chronic neuroinflammation, contributing to the initiation and progression of neurodegenerative diseases such as in AD or Parkinson disease (PD) [107, 108]. Correspondingly, previous reports identified IAV infection as a risk factor for PD or the development of PD-like symptoms [109–111].

Altogether, in this study we are able to demonstrate that respiratory infection with IAV PR8/A/34(H1N1) accumulates in the activation of microglia, dysbalanced glutamatergic neurotransmission and neurotrophin gene expression. Furthermore, we highlight the influence of peripheral inflammation on the immune cell homeostasis of the brain influencing neuronal integrity. Lastly, we provide a novel and highly sensitive approach to characterize the composition of synaptosomes via flow cytometry. Establishing our new method allowed us to gain further insight into the mechanisms involved in the development of neuronal alterations described before, but far more allows an application to various other research domains in a future perspective. We propose Flow Synaptometry as an additional tool, besides immunofluorescence microscopy and histological examination, to quantitatively assess the synapse integrity with high sensitivity in homeostatic conditions and to unravel modest synaptic changes during time-limited neuroinflammatory processes or the onset phases of neuroneurodegenerative conditions.

## 9. Acknowledgments

The authors want to thank Petra Grüneberg, Dr. Sarah Abidat Schneider, Tatjana Hirsch, Kathrin Pohlmann, Sybille Tschorn, and Dr. Christel Bonnas for their excellent technical assistance. We further acknowledge the support of the Combinatorial NeuroImaging Core Facility at the Leibniz Institute of Neurobiology in Magdeburg.

## 10. Authors’ Contributions

HPD, DB, and IRD designed and organized the experiments. HPD, CAF, KS, CE, and HFZ conducted the experiments. HPD, JS, JDB, and HFZ analyzed data. HPD, CAF, JS, MK, HFZ, KHS, DD, AK, DB and IRD interpreted data. HPD, JS, AK, DB and IRD wrote the paper. AK and IRD supervised the study.

## 11. Ethics Approval

The study was performed in accordance with the German National Guidelines for the Use of Experimental Animals and the protocol was approved by the Landesverwaltungsamt Sachsen-Anhalt. Food and water were available *ad libitum*. All efforts were done to minimize the suffering of mice used in this investigation.

## 12. Funding

The work was supported by grants from the German Research Foundation within the CRC854 (Project A25) to MK and IRD and the European Structural and Investment Funds (ESF, ZS/2016/08/80645) to IRD. Moreover, DB received funding from the German Research Foundation (BR2221/6-1).

## 13. Competing Interests

This study used antibodies from Synaptic Systems (Göttingen, Germany) of which CE is CEO. The other authors declare no competing financial interests.

---

**Suppl. Fig. 1:**
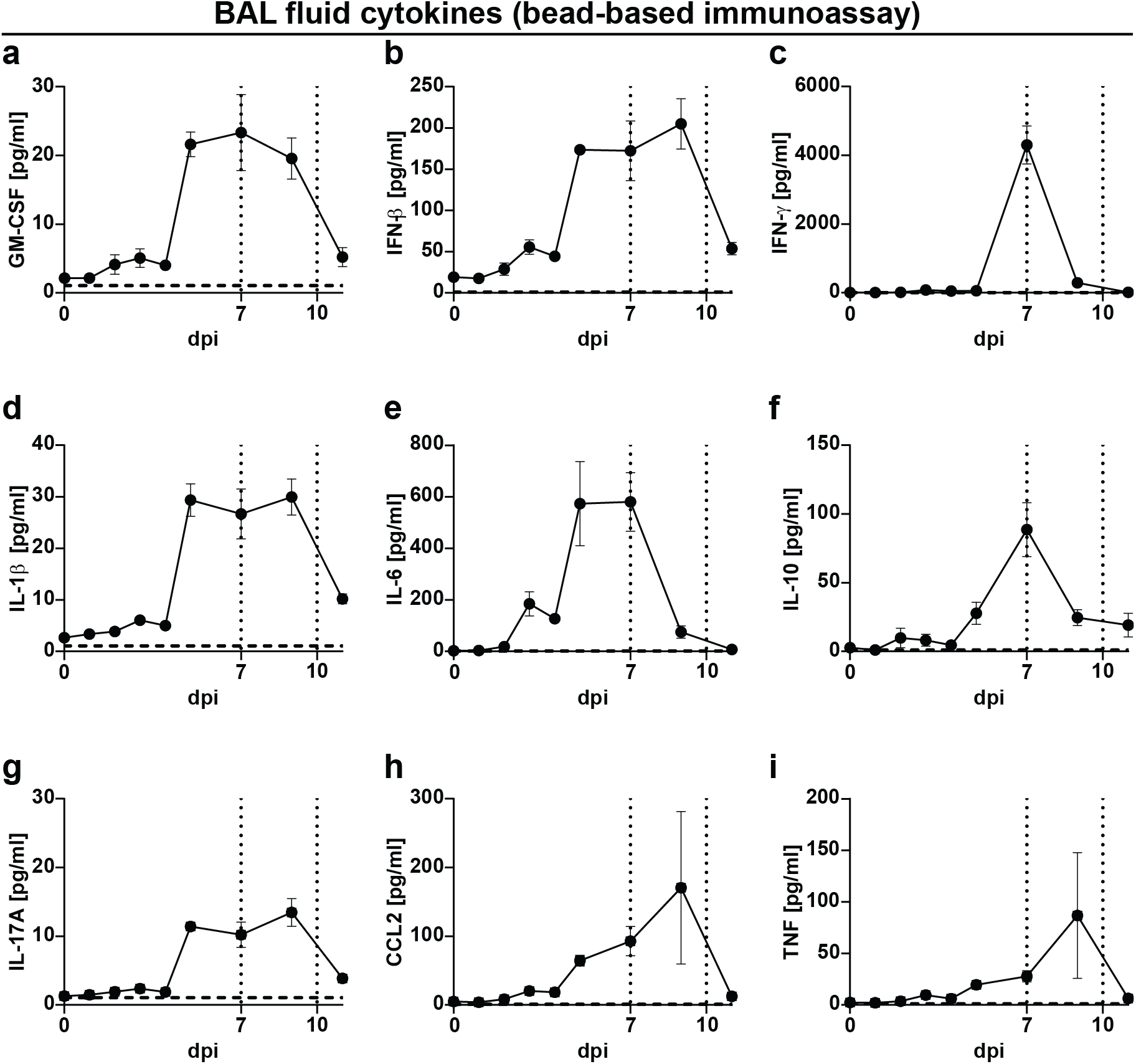
Lung cytokine levels during the course of IAV PR8/A/34(H1N1) infection. (**a-i**) Cytokine levels in the lung of infected mice from 0 to 11 dpi. Dashed vertical lines indicate the time points of experiments in line with the experimental model and dashed horizontal lines indicate the detection limit of each cytokine, respectively. Data are shown as mean ± SEM.

**Suppl. Fig. 2:**
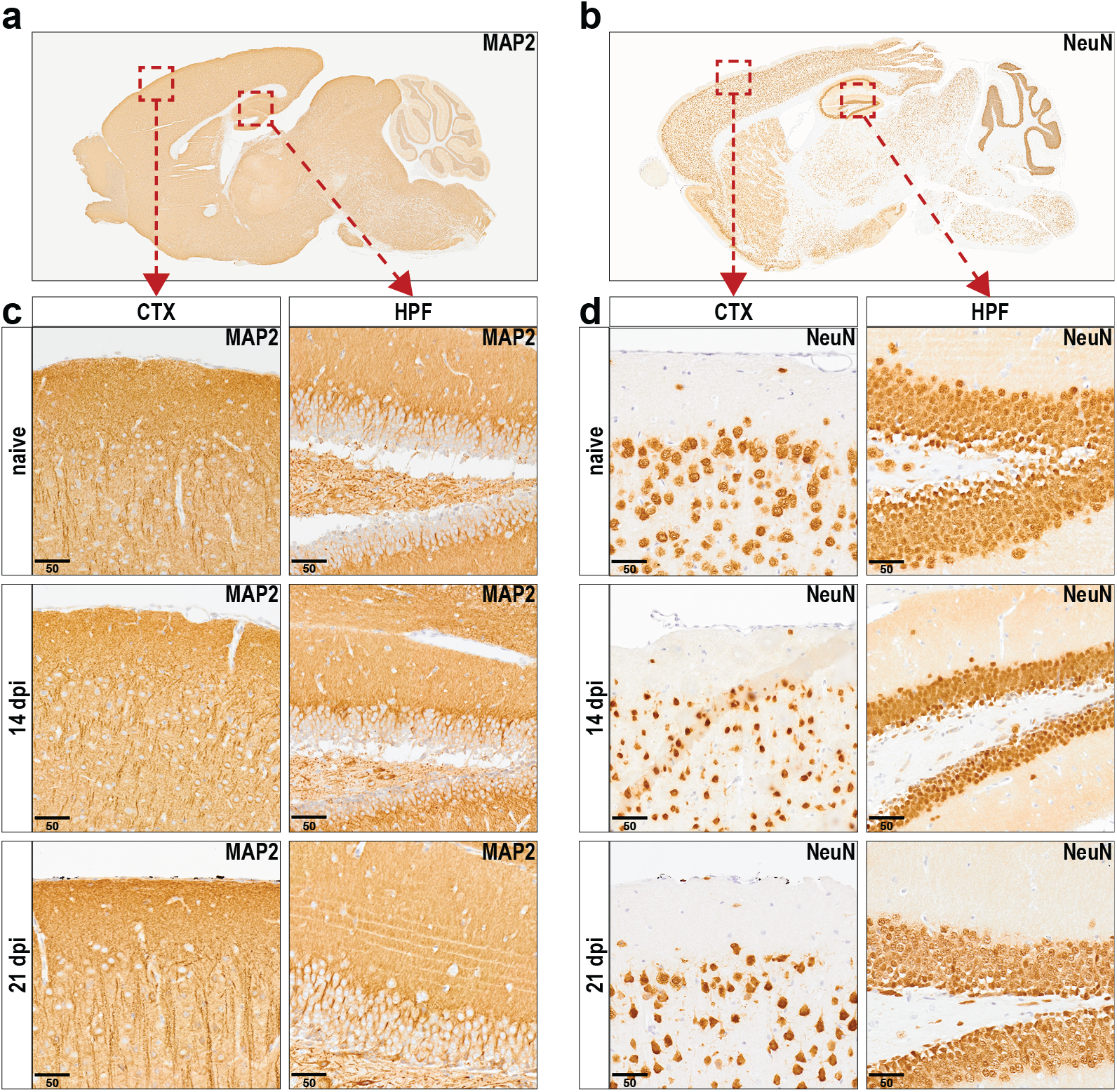
Histopathological examination of neuronal markers in brain tissue upon infection with IAV. (**a, b**) Histopathological overview of representative sagittal paraffin sections from brains of naive mice stained against MAP2 or NeuN. (c, d) Panels show the Cortex (CTX, left panels) and Hippocampal formation region (HPF, right panels) from naive and infected mice (14 and 21 dpi) upon staining against MAP2 or NeuN. Scale bars = 50 µm.

**Suppl. Fig. 3:**
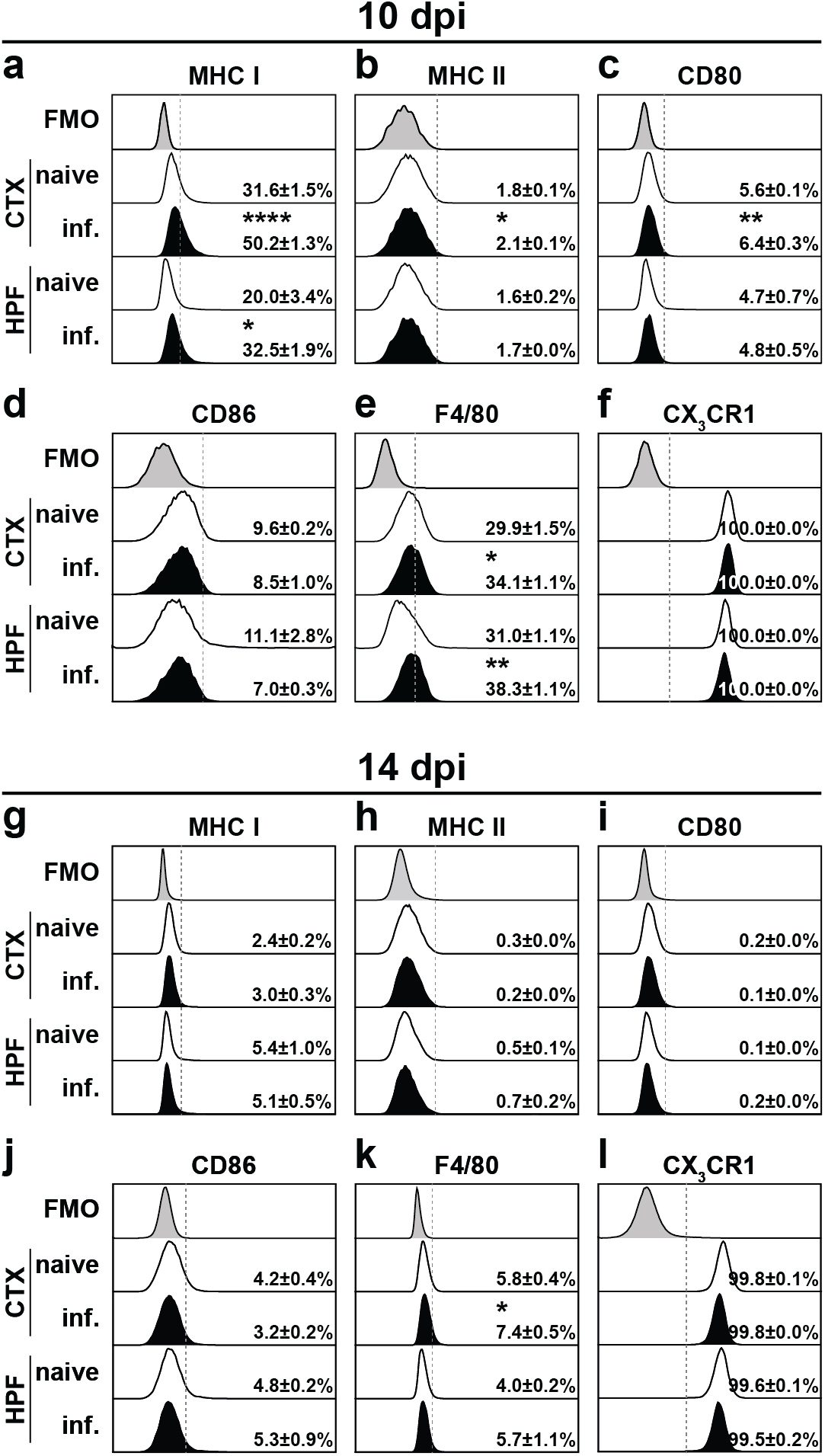
Phenotypical characterization of microglia cells in the late phase of IAV infection. Phenotype of microglia cells was analyzed by flow cytometry. (**a-l**) Histograms show the representative expression level of the surface marker by cells (naive mice without tint, IAV-infected mice tinted) in comparison to the corresponding FMO control (grey tint) at 10 dpi or 14 dpi. Dashed vertical line and numbers mark cells positively expressing the surface marker (± SEM). Groups were compared via Student’s *t*-test with Welch’s correction and significant differences are indicated by * (*p* < .05, ** *p* < .01, *** *p* < .001, **** *p* < .0001).

## Notes

### Competing Interest Statement

This study used antibodies from Synaptic Systems (Goettingen, Germany) of which CE is CEO. The other authors declare no competing financial interests.

## References

1. Kreijtz JHCM, Fouchier RAM, Rimmelzwaan GF (2011) Immune responses to influenza virus infection. Virus Res 162:19–30. https://doi.org/10.1016/j.virusres.2011.09.022

2. Centers for Disease Control and Prevention (2019) Seasonal Influenza (Flu): Background and Epidemiology. https://www.cdc.gov/flu/professionals/acip/background-epidemiology.htm

3. Thompson WW, Weintraub E, Dhankhar P, Cheng P-Y, Brammer L, Meltzer MI, Bresee JS, Shay DK (2009) Estimates of US influenza-associated deaths made using four different methods. Influenza Other Respir Viruses 3:37–49. https://doi.org/10.1111/j.1750-2659.2009.00073.x

4. European Centre for Disease Prevention and Control Factsheet about seasonal influenza. https://www.ecdc.europa.eu/en/seasonal-influenza/facts/factsheet. Accessed 12 Apr 2021

5. European Centre for Disease Prevention and Control (2020) Seasonal influenza 2019-2020. In: ECDC (ed) Annual epidemiological report for 2019, Stockholm

6. Haller O, Staeheli P, Kochs G (2008) Protective role of interferon-induced Mx GTPases against influenza viruses. Revue scientifique et technique (International Office of Epizootics) 28

7. Bahadoran A, Lee SH, Wang SM, Manikam R, Rajarajeswaran J, Raju CS, Sekaran SD (2016) Immune Responses to Influenza Virus and Its Correlation to Age and Inherited Factors. Front Microbiol 7:1841. https://doi.org/10.3389/fmicb.2016.01841

8. Dantzer R (2001) Cytokine-induced sickness behavior: mechanisms and implications. Annals of the New York Academy of Sciences 933:222–234. https://doi.org/10.1111/j.1749-6632.2001.tb05827.x

9. Banks WA, Erickson MA (2010) The blood–brain barrier and immune function and dysfunction. Neurobiology of Disease 37:26–32. https://doi.org/10.1016/j.nbd.2009.07.031

10. Pan W, Stone KP, Hsuchou H, Manda VK, Zhang Y, Kastin AJ (2011) Cytokine signaling modulates blood-brain barrier function. Curr Pharm Des 17:3729–3740. https://doi.org/10.2174/138161211798220918

11. Jang H, Boltz D, McClaren J, Pani AK, Smeyne M, Korff A, Webster R, Smeyne RJ (2012) Inflammatory effects of highly pathogenic H5N1 influenza virus infection in the CNS of mice. J Neurosci 32:1545–1559. https://doi.org/10.1523/JNEUROSCI.5123-11.2012

12. Jurgens HA, Amancherla K, Johnson RW (2012) Influenza infection induces neuroinflammation, alters hippocampal neuron morphology, and impairs cognition in adult mice. The Journal of neuroscience: the official journal of the Society for Neuroscience 32:3958– 3968. https://doi.org/10.1523/JNEUROSCI.6389-11.2012

13. Hosseini S, Wilk E, Michaelsen-Preusse K, Gerhauser I, Baumgärtner W, Geffers R, Schughart K, Korte M (2018) Long-Term Neuroinflammation Induced by Influenza A Virus Infection and the Impact on Hippocampal Neuron Morphology and Function. J Neurosci 38:3060–3080. https://doi.org/10.1523/JNEUROSCI.1740-17.2018

14. Hosseini S, Michaelsen-Preusse K, Schughart K, Korte M (2021) Long-Term Consequence of Non-neurotropic H3N2 Influenza A Virus Infection for the Progression of Alzheimer’s Disease Symptoms. Front Cell Neurosci 15. https://doi.org/10.3389/fncel.2021.643650

15. Nimmerjahn A, Kirchhoff F, Helmchen F (2005) Resting microglial cells are highly dynamic surveillants of brain parenchyma in vivo. Science 308:1314–1318. https://doi.org/10.1126/science.1110647

16. Bar E, Barak B (2019) Microglia roles in synaptic plasticity and myelination in homeostatic conditions and neurodevelopmental disorders. Glia 67:2125–2141. https://doi.org/10.1002/glia.23637

17. Prinz M, Jung S, Priller J (2019) Microglia Biology: One Century of Evolving Concepts. Cell 179:292–311. https://doi.org/10.1016/j.cell.2019.08.053

18. Bilbo S, Stevens B (2017) Microglia: The Brain’s First Responders. Cerebrum 2017

19. Wolf SA, Boddeke HWGM, Kettenmann H (2017) Microglia in Physiology and Disease. Annu Rev Physiol 79:619–643. https://doi.org/10.1146/annurev-physiol-022516-034406

20. Salter MW, Stevens B (2017) Microglia emerge as central players in brain disease. Nat Med 23:1018–1027. https://doi.org/10.1038/nm.4397

21. Li Q, Barres BA (2018) Microglia and macrophages in brain homeostasis and disease. Nat Rev Immunol 18:225–242. https://doi.org/10.1038/nri.2017.125

22. Gibney SM, McGuinness B, Prendergast C, Harkin A, Connor TJ (2013) Poly I:C-induced activation of the immune response is accompanied by depression and anxiety-like behaviours, kynurenine pathway activation and reduced BDNF expression. Brain Behav Immun 28:170–181. https://doi.org/10.1016/j.bbi.2012.11.010

23. Savage JC, St-Pierre M-K, Hui CW, Tremblay M-E (2019) Microglial Ultrastructure in the Hippocampus of a Lipopolysaccharide-Induced Sickness Mouse Model. Front Neurosci 13:1340. https://doi.org/10.3389/fnins.2019.01340

24. Lasselin J, Treadway MT, Lacourt TE, Soop A, Olsson MJ, Karshikoff B, Paues-Göranson S, Axelsson J, Dantzer R, Lekander M (2017) Lipopolysaccharide Alters Motivated Behavior in a Monetary Reward Task: a Randomized Trial. Neuropsychopharmacology 42:801–810. https://doi.org/10.1038/npp.2016.191

25. Stegemann S, Dahlberg S, Kröger A, Gereke M, Bruder D, Henriques-Normark B, Gunzer M, Tailleux L (2009) Increased Susceptibility for Superinfection with Streptococcus pneumoniae during Influenza Virus Infection Is Not Caused by TLR7-Mediated Lymphopenia. PLoS ONE 4:e4840. https://doi.org/10.1371/journal.pone.0004840

26. Stegemann-Koniszewski S, Behrens S, Boehme JD, Hochnadel I, Riese P, Guzmán CA, Kröger A, Schreiber J, Gunzer M, Bruder D (2018) Respiratory Influenza A Virus Infection Triggers Local and Systemic Natural Killer Cell Activation via Toll-Like Receptor 7. Front Immunol 9:245. https://doi.org/10.3389/fimmu.2018.00245

27. Düsedau HP, Kleveman J, Figueiredo CA, Biswas A, Steffen J, Kliche S, Haak S, Za-grebelsky M, Korte M, Dunay IR (2019) p75NTR regulates brain mononuclear cell function and neuronal structure in Toxoplasma infection-induced neuroinflammation. Glia 67:193–211. https://doi.org/10.1002/glia.23553

28. Lein ES, Hawrylycz MJ, Ao N, Ayres M, Bensinger A, Bernard A, Boe AF, Boguski MS, Brockway KS, Byrnes EJ, Chen L, Chen L, Chen T-M, Chin MC, Chong J, Crook BE, Czaplinska A, Dang CN, Datta S, Dee NR, Desaki AL, Desta T, Diep E, Dolbeare TA, Donelan MJ, Dong H-W, Dougherty JG, Duncan BJ, Ebbert AJ, Eichele G, Estin LK, Faber C, Facer BA, Fields R, Fischer SR, Fliss TP, Frensley C, Gates SN, Glattfelder KJ, Halverson KR, Hart MR, Hohmann JG, Howell MP, Jeung DP, Johnson RA, Karr PT, Kawal R, Kidney JM, Knapik RH, Kuan CL, Lake JH, Laramee AR, Larsen KD, Lau C, Lemon TA, Liang AJ, Liu Y, Luong LT, Michaels J, Morgan JJ, Morgan RJ, Mortrud MT, Mosqueda NF, Ng LL, Ng R, Orta GJ, Overly CC, Pak TH, Parry SE, Pathak SD, Pearson OC, Puchalski RB, Riley ZL, Rockett HR, Rowland SA, Royall JJ, Ruiz MJ, Sarno NR, Schaffnit K, Shapovalova NV, Sivisay T, Slaughterbeck CR, Smith SC, Smith KA, Smith BI, Sodt AJ, Stewart NN, Stumpf K-R, Sunkin SM, Sutram M, Tam A, Teemer CD, Thaller C, Thompson CL, Varnam LR, Visel A, Whitlock RM, Wohnoutka PE, Wolkey CK, Wong VY, Wood M, Yaylaoglu MB, Young RC, Youngstrom BL, Yuan XF, Zhang B, Zwing-man TA, Jones AR (2007) Genome-wide atlas of gene expression in the adult mouse brain. Nature 445:168–176. https://doi.org/10.1038/nature05453

29. Figueiredo CA, Düsedau HP, Steffen J, Gupta N, Dunay MP, Toth GK, Reglodi D, Heimesaat MM, Dunay IR (2019) Immunomodulatory Effects of the Neuropeptide Pituitary Adenylate Cyclase-Activating Polypeptide in Acute Toxoplasmosis. Front Cell Infect Microbiol 9:154. https://doi.org/10.3389/fcimb.2019.00154

30. Sharma-Chawla N, Sender V, Kershaw O, Gruber AD, Volckmar J, Henriques-Normark B, Stegemann-Koniszewski S, Bruder D (2016) Influenza A Virus Infection Predisposes Hosts to Secondary Infection with Different Streptococcus pneumoniae Serotypes with Similar Outcome but Serotype-Specific Manifestation. Infect Immun 84:3445–3457. https://doi.org/10.1128/IAI.00422-16

31. Schindelin J, Arganda-Carreras I, Frise E, Kaynig V, Longair M, Pietzsch T, Preibisch S, Rueden C, Saalfeld S, Schmid B, Tinevez J-Y, White DJ, Hartenstein V, Eliceiri K, To-mancak P, Cardona A (2012) Fiji: An open-source platform for biological-image analysis. Nat Methods 9:676–682. https://doi.org/10.1038/nmeth.2019

32. Smalla K-H, Klemmer P, Wyneken U (2013) Isolation of the Postsynaptic Density: A Specialization of the Subsynaptic Cytoskeleton. In: Dermietzel R (ed) The cytoskeleton: Imaging, isolation, and interaction, vol 79. Humana Press, New York, pp 265–280

33. Hobson BD, Sims PA (2019) Critical Analysis of Particle Detection Artifacts in Synaptosome Flow Cytometry. eNeuro 6. https://doi.org/10.1523/ENEURO.0009-19.2019

34. Breukel AI, Besselsen E, Ghijsen WE (1997) Synaptosomes. A model system to study release of multiple classes of neurotransmitters. Methods Mol Biol 72:33–47. https://doi.org/10.1385/0-89603-394-5:33

35. R Core Team (2020) R: A language and environment for statistical computing. R Foundation for Statistical Computing, Vienna, Austria

36. Sarkar D (2008) Lattice: Multivariate Data Visualization with R. Springer New York, New York, NY

37. Layé S, Parnet P, Goujon E, Dantzer R (1994) Peripheral administration of lipopolysaccharide induces the expression of cytokine transcripts in the brain and pituitary of mice. Molecular Brain Research 27:157–162. https://doi.org/10.1016/0169-328X(94)90197-X

38. Blank T, Detje CN, Spieß A, Hagemeyer N, Brendecke SM, Wolfart J, Staszewski O, Zöller T, Papageorgiou I, Schneider J, Paricio-Montesinos R, Eisel ULM, Manahan-Vaughan D, Jansen S, Lienenklaus S, Lu B, Imai Y, Müller M, Goelz SE, Baker DP, Schwaninger M, Kann O, Heikenwalder M, Kalinke U, Prinz M (2016) Brain Endothelial- and Epithelial-Specific Interferon Receptor Chain 1 Drives Virus-Induced Sickness Behavior and Cognitive Impairment. Immunity 44:901–912. https://doi.org/10.1016/j.immuni.2016.04.005

39. Daniels BP, Holman DW, Cruz-Orengo L, Jujjavarapu H, Durrant DM, Klein RS (2014) Viral pathogen-associated molecular patterns regulate blood-brain barrier integrity via competing innate cytokine signals. mBio 5:e01476–14. https://doi.org/10.1128/mBio.01476-14

40. Dyrna F, Hanske S, Krueger M, Bechmann I (2013) The blood-brain barrier. J Neuroimmune Pharmacol 8:763–773. https://doi.org/10.1007/s11481-013-9473-5

41. Klein RS, Lin E, Zhang B, Luster AD, Tollett J, Samuel MA, Engle M, Diamond MS (2005) Neuronal CXCL10 directs CD8+ T-cell recruitment and control of West Nile virus encephalitis. Journal of Virology 79:11457–11466. https://doi.org/10.1128/JVI.79.17.11457-11466.2005

42. Zhang B, Chan YK, Lu B, Diamond MS, Klein RS (2008) CXCR3 mediates region-specific antiviral T cell trafficking within the central nervous system during West Nile virus encephalitis. J Immunol 180:2641–2649. https://doi.org/10.4049/jimmunol.180.4.2641

43. Kallfass C, Ackerman A, Lienenklaus S, Weiss S, Heimrich B, Staeheli P (2012) Visualizing production of beta interferon by astrocytes and microglia in brain of La Crosse virus-infected mice. Journal of Virology 86:11223–11230. https://doi.org/10.1128/JVI.0109312

44. Owens T, Khorooshi R, Wlodarczyk A, Asgari N (2014) Interferons in the central nervous system: a few instruments play many tunes. Glia 62:339–355. https://doi.org/10.1002/glia.22608

45. Bennett ML, Bennett FC, Liddelow SA, Ajami B, Zamanian JL, Fernhoff NB, Mulinyawe SB, Bohlen CJ, Adil A, Tucker A, Weissman IL, Chang EF, Li G, Grant GA, Hayden Gephart MG, Barres BA (2016) New tools for studying microglia in the mouse and human CNS. Proc Natl Acad Sci U S A 113:E1738–46. https://doi.org/10.1073/pnas.1525528113

46. Ransohoff RM (2016) A polarizing question: do M1 and M2 microglia exist? Nat Neurosci 19:987–991. https://doi.org/10.1038/nn.4338

47. Dubbelaar ML, Kracht L, Eggen BJL, Boddeke EWGM (2018) The Kaleidoscope of Microglial Phenotypes. Front Immunol 9:1753. https://doi.org/10.3389/fimmu.2018.01753

48. Kettenmann H, Kirchhoff F, Verkhratsky A (2013) Microglia: new roles for the synaptic stripper. Neuron 77:10–18. https://doi.org/10.1016/j.neuron.2012.12.023

49. Brown GC, Neher JJ (2014) Microglial phagocytosis of live neurons. Nat Rev Neurosci 15:209–216. https://doi.org/10.1038/nrn3710

50. Kim H-J, Cho M-H, Shim WH, Kim JK, Jeon E-Y, Kim D-H, Yoon S-Y (2017) Deficient autophagy in microglia impairs synaptic pruning and causes social behavioral defects. Mol Psychiatry 22:1576–1584. https://doi.org/10.1038/mp.2016.103

51. Neniskyte U, Gross CT (2017) Errant gardeners: glial-cell-dependent synaptic pruning and neurodevelopmental disorders. Nat Rev Neurosci 18:658–670. https://doi.org/10.1038/nrn.2017.110

52. Sellgren CM, Gracias J, Watmuff B, Biag JD, Thanos JM, Whittredge PB, Fu T, Worringer K, Brown HE, Wang J, Kaykas A, Karmacharya R, Goold CP, Sheridan SD, Perlis RH (2019) Increased synapse elimination by microglia in schizophrenia patient-derived models of synaptic pruning. Nat Neurosci 22:374–385. https://doi.org/10.1038/s41593-018-0334-7

53. Stevens B, Allen NJ, Vazquez LE, Howell GR, Christopherson KS, Nouri N, Micheva KD, Mehalow AK, Huberman AD, Stafford B, Sher A, Litke AM, Lambris JD, Smith SJ, John SWM, Barres BA (2007) The classical complement cascade mediates CNS synapse elimination. Cell 131:1164–1178. https://doi.org/10.1016/j.cell.2007.10.036

54. Kober DL, Brett TJ (2017) TREM2-Ligand Interactions in Health and Disease. J Mol Biol 429:1607–1629. https://doi.org/10.1016/j.jmb.2017.04.004

55. Lang D, Schott BH, van Ham M, Morton L, Kulikovskaja L, Herrera-Molina R, Pielot R, Klawonn F, Montag D, Jänsch L, Gundelfinger ED, Smalla KH, Dunay IR (2018) Chronic Toxoplasma infection is associated with distinct alterations in the synaptic protein composition. J Neuroinflammation 15:438. https://doi.org/10.1186/s12974-018-1242-1

56. Biesemann C, Grønborg M, Luquet E, Wichert SP, Bernard V, Bungers SR, Cooper B, Varoqueaux F, Li L, Byrne JA, Urlaub H, Jahn O, Brose N, Herzog E (2014) Proteomic screening of glutamatergic mouse brain synaptosomes isolated by fluorescence activated sorting. EMBO J 33:157–170. https://doi.org/10.1002/embj.201386120

57. Gylys KH, Bilousova T (2017) Flow Cytometry Analysis and Quantitative Characterization of Tau in Synaptosomes from Alzheimer’s Disease Brains. Methods Mol Biol 1523:273–284. https://doi.org/10.1007/978-1-4939-6598-4_16

58. Gray EG, Whittaker VP (1962) The isolation of nerve endings from brain: an electronmicroscopic study of cell fragments derived by homogenization and centrifugation. J Anatomy 96:79–88

59. Dunkley PR, Jarvie PE, Heath JW, Kidd GJ, Rostas JA (1986) A rapid method for isolation of synaptosomes on Percoll gradients. Brain Research 372:115–129. https://doi.org/10.1016/0006-8993(86)91464-2

60. Chao MV (2003) Neurotrophins and their receptors: a convergence point for many signalling pathways. Nat Rev Neurosci 4:299–309. https://doi.org/10.1038/nrn1078

61. Cunha C, Brambilla R, Thomas KL (2010) A simple role for BDNF in learning and memory? Front Mol Neurosci 3:1. https://doi.org/10.3389/neuro.02.001.2010

62. Meeker R, Williams K (2014) Dynamic nature of the p75 neurotrophin receptor in response to injury and disease. J Neuroimmune Pharmacol 9:615–628. https://doi.org/10.1007/s11481-014-9566-9

63. Ibáñez CF, Simi A (2012) p75 neurotrophin receptor signaling in nervous system injury and degeneration: Paradox and opportunity. Trends Neurosci 35:431–440. https://doi.org/10.1016/j.tins.2012.03.007

64. Müller N, Schwarz MJ (2007) The immune-mediated alteration of serotonin and glutamate: towards an integrated view of depression. Mol Psychiatry 12:988–1000. https://doi.org/10.1038/sj.mp.4002006

65. Zhang XZ, Penzel T, Han F (2013) Increased incidence of narcolepsy following the 2009 H1N1 pandemic. Somnologie 17:90–93. https://doi.org/10.1007/s11818-013-0619-8

66. Bornand D, Toovey S, Jick SS, Meier CR (2016) The risk of new onset depression in association with influenza--A population-based observational study. Brain Behav Immun 53:131–137. https://doi.org/10.1016/j.bbi.2015.12.005

67. Cai KC, van Mil S, Murray E, Mallet J-F, Matar C, Ismail N (2016) Age and sex differences in immune response following LPS treatment in mice. Brain Behav Immun 58:327– 337. https://doi.org/10.1016/j.bbi.2016.08.002

68. Gandhi R, Hayley S, Gibb J, Merali Z, Anisman H (2007) Influence of poly I:C on sickness behaviors, plasma cytokines, corticosterone and central monoamine activity: moderation by social stressors. Brain Behav Immun 21:477–489. https://doi.org/10.1016/j.bbi.2006.12.005

69. Dantzer R, O’Connor JC, Freund GG, Johnson RW, Kelley KW (2008) From inflammation to sickness and depression: when the immune system subjugates the brain. Nat Rev Neurosci 9:46–56. https://doi.org/10.1038/nrn2297

70. Kraus J, Voigt K, Schuller AM, Scholz M, Kim KS, Schilling M, Schäbitz WR, Oschmann P, Engelhardt B (2008) Interferon-beta stabilizes barrier characteristics of the blood-brain barrier in four different species in vitro. Mult Scler 14:843–852. https://doi.org/10.1177/1352458508088940

71. Veldhuis WB, Floris S, van der Meide PH, Vos IMP, Vries HE de, Dijkstra CD, Bär PR, Nicolay K (2003) Interferon-beta prevents cytokine-induced neutrophil infiltration and attenuates blood-brain barrier disruption. J Cereb Blood Flow Metab 23:1060–1069. https://doi.org/10.1097/01.WCB.0000080701.47016.24

72. Chai Q, He WQ, Zhou M, Lu H, Fu ZF (2014) Enhancement of blood-brain barrier permeability and reduction of tight junction protein expression are modulated by chemo-kines/cytokines induced by rabies virus infection. Journal of Virology 88:4698–4710. https://doi.org/10.1128/JVI.03149-13

73. Utech M, Ivanov AI, Samarin SN, Bruewer M, Turner JR, Mrsny RJ, Parkos CA, Nusrat A (2005) Mechanism of IFN-gamma-induced endocytosis of tight junction proteins: myo-sin II-dependent vacuolarization of the apical plasma membrane. Mol Biol Cell 16:5040– 5052. https://doi.org/10.1091/mbc.e05-03-0193

74. Monteiro S, Roque S, Marques F, Correia-Neves M, Cerqueira JJ (2017) Brain interference: Revisiting the role of IFNγ in the central nervous system. Progress in Neurobiology 156:149–163. https://doi.org/10.1016/j.pneurobio.2017.05.003

75. Loos T, Dekeyzer L, Struyf S, Schutyser E, Gijsbers K, Gouwy M, Fraeyman A, Put W, Ronsse I, Grillet B, Opdenakker G, van Damme J, Proost P (2006) TLR ligands and cytokines induce CXCR3 ligands in endothelial cells: enhanced CXCL9 in autoimmune arthritis. Lab Invest 86:902–916. https://doi.org/10.1038/labinvest.3700453

76. Gack MU, Albrecht RA, Urano T, Inn K-S, Huang I-C, Carnero E, Farzan M, Inoue S, Jung JU, García-Sastre A (2009) Influenza A virus NS1 targets the ubiquitin ligase TRIM25 to evade recognition by the host viral RNA sensor RIG-I. Cell Host Microbe 5:439–449. https://doi.org/10.1016/j.chom.2009.04.006

77. Chung W-C, Kang H-R, Yoon H, Kang S-J, Ting JP-Y, Song MJ (2015) Influenza A Virus NS1 Protein Inhibits the NLRP3 Inflammasome. PLoS ONE 10:e0126456. https://doi.org/10.1371/journal.pone.0126456

78. Oishi K, Yamayoshi S, Kawaoka Y (2015) Mapping of a Region of the PA-X Protein of Influenza A Virus That Is Important for Its Shutoff Activity. Journal of Virology 89:8661– 8665. https://doi.org/10.1128/JVI.01132-15

79. Vasek MJ, Garber C, Dorsey D, Durrant DM, Bollman B, Soung A, Yu J, Perez-Torres C, Frouin A, Wilton DK, Funk K, DeMasters BK, Jiang X, Bowen JR, Mennerick S, Robinson JK, Garbow JR, Tyler KL, Suthar MS, Schmidt RE, Stevens B, Klein RS (2016) A complement-microglial axis drives synapse loss during virus-induced memory impairment. Nature 534:538–543. https://doi.org/10.1038/nature18283

80. French T, Düsedau HP, Steffen J, Biswas A, Ahmed N, Hartmann S, Schüler T, Schott BH, Dunay IR (2019) Neuronal impairment following chronic Toxoplasma gondii infection is aggravated by intestinal nematode challenge in an IFN-γ-dependent manner. J Neuroinflammation 16:159. https://doi.org/10.1186/s12974-019-1539-8

81. Postupna NO, Keene CD, Latimer C, Sherfield EE, van Gelder RD, Ojemann JG, Montine TJ, Darvas M (2014) Flow cytometry analysis of synaptosomes from post-mortem human brain reveals changes specific to Lewy body and Alzheimer’s disease. Lab Invest 94:1161– 1172. https://doi.org/10.1038/labinvest.2014.103

82. Postupna NO, Latimer CS, Keene CD, Montine KS, Montine TJ, Darvas M (2018) Flow cytometric evaluation of crude synaptosome preparation as a way to study synaptic alteration in neurodegenerative diseases. Neuromethods 141:297–310. https://doi.org/10.1007/978-1-4939-8739-9_17

83. Gajera CR, Fernandez R, Montine KS, Fox EJ, Mrdjen D, Postupna NO, Keene CD, Bendall SC, Montine TJ (2021) Mass-tag barcoding for multiplexed analysis of human synaptosomes and other anuclear events. Cytometry A. https://doi.org/10.1002/cyto.a.24340

84. Hunsberger HC, Wang D, Petrisko TJ, Alhowail A, Setti SE, Suppiramaniam V, Konat GW, Reed MN (2016) Peripherally restricted viral challenge elevates extracellular gluta-mate and enhances synaptic transmission in the hippocampus. J Neurochem 138:307–316. https://doi.org/10.1111/jnc.13665

85. Stone TW, Darlington LG (2002) Endogenous kynurenines as targets for drug discovery and development. Nat Rev Drug Discov 1:609–620. https://doi.org/10.1038/nrd870

86. Jeon SW, Kim Y-K (2017) Inflammation-induced depression: Its pathophysiology and therapeutic implications. J Neuroimmunol 313:92–98. https://doi.org/10.1016/j.jneuroim.2017.10.016

87. Carlin JM, Borden EC, Sondel PM, Byrne GI (1989) Interferon-induced indoleamine 2,3-dioxygenase activity in human mononuclear phagocytes. J Leukoc Biol 45:29–34. https://doi.org/10.1002/jlb.45.1.29

88. Fujigaki H, Saito K, Fujigaki S, Takemura M, Sudo K, Ishiguro H, Seishima M (2006) The signal transducer and activator of transcription 1alpha and interferon regulatory factor 1 are not essential for the induction of indoleamine 2,3-dioxygenase by lipopolysaccharide: involvement of p38 mitogen-activated protein kinase and nuclear factor-kappaB pathways, and synergistic effect of several proinflammatory cytokines. J Biochem 139:655–662. https://doi.org/10.1093/jb/mvj072

89. Zunszain PA, Anacker C, Cattaneo A, Choudhury S, Musaelyan K, Myint AM, Thuret S, Price J, Pariante CM (2012) Interleukin-1β: a new regulator of the kynurenine pathway affecting human hippocampal neurogenesis. Neuropsychopharmacology 37:939–949. https://doi.org/10.1038/npp.2011.277

90. Gaelings L, Söderholm S, Bugai A, Fu Y, Nandania J, Schepens B, Lorey MB, Tynell J, Vande Ginste L, Le Goffic R, Miller MS, Kuisma M, Marjomäki V, Brabander J de, Matikainen S, Nyman TA, Bamford DH, Saelens X, Julkunen I, Paavilainen H, Hukkanen V, Velagapudi V, Kainov DE (2017) Regulation of kynurenine biosynthesis during influenza virus infection. FEBS J 284:222–236. https://doi.org/10.1111/febs.13966

91. Baumgartner R, Forteza MJ, Ketelhuth DFJ (2019) The interplay between cytokines and the Kynurenine pathway in inflammation and atherosclerosis. Cytokine 122:154148. https://doi.org/10.1016/j.cyto.2017.09.004

92. Suh H-S, Zhao M-L, Rivieccio M, Choi S, Connolly E, Zhao Y, Takikawa O, Brosnan CF, Lee SC (2007) Astrocyte indoleamine 2,3-dioxygenase is induced by the TLR3 ligand poly(I:C): mechanism of induction and role in antiviral response. Journal of Virology 81:9838–9850. https://doi.org/10.1128/JVI.00792-07

93. Ciranna L (2006) Serotonin as a modulator of glutamate- and GABA-mediated neurotransmission: implications in physiological functions and in pathology. Curr Neuropharmacol 4:101–114. https://doi.org/10.2174/157015906776359540

94. O’Connor JC, Lawson MA, André C, Moreau M, Lestage J, Castanon N, Kelley KW, Dantzer R (2009) Lipopolysaccharide-induced depressive-like behavior is mediated by indoleamine 2,3-dioxygenase activation in mice. Mol Psychiatry 14:511–522. https://doi.org/10.1038/sj.mp.4002148

95. Schwarcz R, Stone TW (2017) The kynurenine pathway and the brain: Challenges, controversies and promises. Neuropharmacology 112:237–247. https://doi.org/10.1016/j.neuropharm.2016.08.003

96. Walker AK, Budac DP, Bisulco S, Lee AW, Smith RA, Beenders B, Kelley KW, Dantzer R (2013) NMDA receptor blockade by ketamine abrogates lipopolysaccharide-induced depressive-like behavior in C57BL/6J mice. Neuropsychopharmacology 38:1609–1616. https://doi.org/10.1038/npp.2013.71

97. Sadasivan S, Zanin M, O’Brien K, Schultz-Cherry S, Smeyne RJ (2015) Induction of microglia activation after infection with the non-neurotropic A/CA/04/2009 H1N1 influenza virus. PLoS ONE 10:e0124047. https://doi.org/10.1371/journal.pone.0124047

98. Guan Z, Fang J (2006) Peripheral immune activation by lipopolysaccharide decreases neurotrophins in the cortex and hippocampus in rats. Brain Behav Immun 20:64–71. https://doi.org/10.1016/j.bbi.2005.04.005

99. Martinowich K, Manji H, Lu B (2007) New insights into BDNF function in depression and anxiety. Nat Neurosci 10:1089–1093. https://doi.org/10.1038/nn1971

100. Leßmann V (1998) Neurotrophin-Dependent Modulation of Glutamatergic Synaptic Transmission in the Mammalian CNS. General Pharmacology: The Vascular System 31:667–674. https://doi.org/10.1016/S0306-3623(98)00190-6

101. Müller N, Schwarz M (2006) Schizophrenia as an inflammation-mediated dysbalance of glutamatergic neurotransmission. Neurotox Res 10:131–148. https://doi.org/10.1007/BF03033242

102. Blaylock RL, Strunecka A (2009) Immune-glutamatergic dysfunction as a central mechanism of the autism spectrum disorders. Curr Med Chem 16:157–170. https://doi.org/10.2174/092986709787002745

103. Ting JT, Feng G (2008) Glutamatergic Synaptic Dysfunction and Obsessive-Compulsive Disorder. Curr Chem Genomics 2:62–75. https://doi.org/10.2174/1875397300802010062

104. Evans DL, Charney DS, Lewis L, Golden RN, Gorman JM, Krishnan KRR, Nemeroff CB, Bremner JD, Carney RM, Coyne JC, Delong MR, Frasure-Smith N, Glassman AH, Gold PW, Grant I, Gwyther L, Ironson G, Johnson RL, Kanner AM, Katon WJ, Kaufmann PG, Keefe FJ, Ketter T, Laughren TP, Leserman J, Lyketsos CG, McDonald WM, McEwen BS, Miller AH, Musselman D, O’Connor C, Petitto JM, Pollock BG, Robinson RG, Roose SP, Rowland J, Sheline Y, Sheps DS, Simon G, Spiegel D, Stunkard A, Sunderland T, Tibbits P, Valvo WJ (2005) Mood disorders in the medically ill: scientific review and recommendations. Biol Psychiatry 58:175–189. https://doi.org/10.1016/j.biopsych.2005.05.001

105. Hosp JA, Dressing A, Blazhenets G, Bormann T, Rau A, Schwabenland M, Thurow J, Wagner D, Waller C, Niesen WD, Frings L, Urbach H, Prinz M, Weiller C, Schroeter N, Meyer PT (2021) Cognitive impairment and altered cerebral glucose metabolism in the subacute stage of COVID-19. Brain. https://doi.org/10.1093/brain/awab009

106. Haley MJ, Brough D, Quintin J, Allan SM (2019) Microglial Priming as Trained Immunity in the Brain. Neuroscience 405:47–54. https://doi.org/10.1016/j.neuroscience.2017.12.039

107. Kempuraj D, Thangavel R, Natteru PA, Selvakumar GP, Saeed D, Zahoor H, Zaheer S, Iyer SS, Zaheer A (2016) Neuroinflammation Induces Neurodegeneration. J Neurol Neurosurg Spine 1

108. Ransohoff RM (2016) How neuroinflammation contributes to neurodegeneration. Science 353:777–783. https://doi.org/10.1126/science.aag2590

109. Sadasivan S, Sharp B, Schultz-Cherry S, Smeyne RJ (2017) Synergistic effects of influenza and 1-methyl-4-phenyl-1,2,3,6-tetrahydropyridine (MPTP) can be eliminated by the use of influenza therapeutics: experimental evidence for the multi-hit hypothesis. NPJ Parkinsons Dis 3:18. https://doi.org/10.1038/s41531-017-0019-z

110. Toovey S, Jick SS, Meier CR (2011) Parkinson’s disease or Parkinson symptoms following seasonal influenza. Influenza Other Respir Viruses 5:328–333. https://doi.org/10.1111/j.1750-2659.2011.00232.x

111. Tulisiak CT, Mercado G, Peelaerts W, Brundin L, Brundin P (2019) Can infections trigger alpha-synucleinopathies? Prog Mol Biol Transl Sci 168:299–322. https://doi.org/10.1016/bs.pmbts.2019.06.002

